# “Towards Passive Dietary Monitoring: Dilated CNN-Based Meal Detection Using Ambulatory High-Resolution Electrogastrography”

**DOI:** 10.64898/2026.06.13.731982

**Authors:** Belinda Chen, Sandya Subramanian

## Abstract

**Background:** Objective tools for longitudinal dietary monitoring, despite its importance in the health triad of diet, sleep, and exercise, remain limited. Widely available ambulatory options include continuous glucose monitoring, which is a delayed response, and manual logging, which is rife with human error. Clinical tools such as gastric emptying scintigraphy are impractical for everyday use. High-resolution electrogastrography (HR-EGG) offers an alternative by treating gastric myoelectric activity as a biomarker of digestive state. However, its utility for ambulatory meal detection remains unclear. We hypothesize that HR-EGG and accelerometry together encode distinct postprandial gastric signatures to enable automated meal detection, and that postural context represents a relevant source of variation in signal detectability.

**Methods:** HR-EGG and accelerometry data were collected from seven healthy adults across sixteen 150-minute meal sessions under IRB-approved protocol. Each session included a 30-minute fasted baseline, consumption of a standardized meal, and 90-minute postprandial period of sitting, walking, and lying in a randomized order. Features of the gastric slow wave, including raw and normalized bandpower, phase gradient directionality (PGD), wave direction, and wave speed, were extracted alongside triaxial accelerometer magnitude. A dilated one-dimensional convolutional network (1D CNN) was trained to classify meal consumption at five-minute resolution using leave-one-subject-out cross-validation. Postural effects on gastric myoelectric metrics were assessed using the Friedman test.

**Results:** The model achieved a mean AUROC of 0.925 (95% CI: [0.857, 0.993]) and mean AUPRC of 0.824 (95% CI: [0.668, 0.980]; null model: 0.20). Feature ablation showed PGD as the most informative input (ΔAUPRC = −0.188), with wave propagation speed the least informative (ΔAUPRC = −0.105). Walking produced the highest signal-to-noise ratio (9.94 dB), lying had the most stable gastric rhythm (89.1% normogastric), and sitting demonstrated the greatest frequency instability (dominant frequency standard deviation = 0.825 cpm).

**Conclusion:** A dilated 1D CNN applied to spatiotemporal HR-EGG features enables temporally aware passive meal detection across ambulatory contexts. This study framework addresses a gap between clinical need for objective dietary monitoring and the limitations of current detection methods.

## Background

Despite its central role in metabolic and gastrointestinal health, eating remains one of the few physiological processes that cannot be monitored objectively and continuously in everyday life. Wearable devices now passively track parameters such as heart rate, physical activity, and sleep, but dietary intake monitoring still relies almost entirely on self-reporting [1]. This gap matters across both clinical and general populations: for the 40 million adults in the United States living with diabetes, meal timing is a crucial component of glycemic management [2]. In gastrointestinal motility disorders such as gastroparesis and functional dyspepsia, the relationship between food intake and symptom onset is also critical to diagnosis and management [3], [4], [5]. Beyond these clinical contexts, passive dietary monitoring could provide the general population with the same objective, low-burden behavioral monitoring that wearable devices already offer for physical activity, heart rate, and sleep. The ideal signal for automated meal detection is one that responds immediately to eating by reflecting the body’s internal physiological response to food, providing a fast and objective confirmation that some caloric content has been consumed.

Current approaches fall short of this ideal in distinct ways. Manual logging through diaries or smartphone applications remains the standard in the clinic and for research studies but is limited by poor long-term compliance and recall bias [6]. Up to 70% of adults misreport their energy intake, and the knowledge that the reported intake will be scrutinized may change eating behaviors [7], [8]. Automated wearable systems instead infer eating through behavioral mechanics, such as acoustic detection of chewing, jaw-mounted sensors, and wrist-worn devices that recognize hand-to-mouth gestures [9], [10], [11], [12]. While reasonably accurate under controlled conditions, these methods are vulnerable to real-life confounds such as talking, drinking, and individual and cultural variations in eating style and meal content, and they focus on the act of eating itself instead of the physiological response. Continuous glucose monitors (CGMs), commonly used in diabetes management, can be used to infer meals from postprandial increases in blood glucose levels, a downstream metabolic response to ingestion [13]. However, the glucose response is delayed relative to the meal depending on macronutrient composition, making it difficult to accurately reconstruct meal timing from the glucose curve alone [13]. In addition, CGMs require a sensor needle inserted under the skin, are disposable and therefore incur cost over time, and are less reliable in individuals with poorly controlled diabetes, where blood glucose levels are influenced by factors beyond recent food intake [14], [15]. Among clinical tools, gastric emptying scintigraphy (GES) remains the gold standard but is resource-intensive, confined to the clinic, and correlates only weakly with symptom severity [4], [16], [17]. Across these methods, none provides a passive, non-invasive readout of the body’s immediate physiological response to a meal in real-world settings.

Stomach muscle signals, also called gastric myoelectric activity, represent a promising alternative, as the stomach generates electrical signals that change in response to food intake. These signals can be measured non-invasively at the body’s surface and provide a direct physiological marker of meal consumption. Gastric myoelectric activity originates in a population of pacemaker cells, the interstitial cells of Cajal (ICC), which produce spontaneous depolarizations known as gastric slow waves at approximately 3 cycles per minute (cpm), or 0.05 Hz [18], [19], [20]. After a meal, the vagus nerve, the primary autonomic pathway linking the brain and the gut, increases its signaling to the stomach. As a result, the amplitude and spectral power of the gastric slow wave increases in a postprandial response that ultimately drives the muscular contractions of digestion [18], [20]. The electrical activity in the stomach muscle is detectable at the abdominal surface and forms the physiological basis for cutaneous electrogastrography (EGG), a non-invasive proxy for digestive state. Traditional EGG, however, uses only one or a few channels and therefore can only capture the temporal frequency of gastric activity [21], [22]. High-resolution electrogastrography (HR-EGG) instead uses dense electrode arrays to resolve the spatial propagation of slow waves across the stomach, including the instantaneous propagation direction, velocity, and phase [23], [24], [25]. Despite these capabilities, the application of HR-EGG to automated meal detection remains understudied.

One of the most critically understudied challenges in ambulatory gastric monitoring is the effect of posture and physical activity on gastric myoelectric signals. Clinical guidelines for electrogastrography (EGG) instruct patients to remain still during recordings because changes in body position can alter signal amplitude and other measured parameters [21], [23], [24]. This sensitivity to position is rooted in autonomic physiology, since as posture changes, it modifies sympathovagal balance which in turn can influence gastric slow-wave activity and motility [26], [27], [28]. Posture may also affect signal quality through mechanical factors like altering the distance between the electrodes and stomach and the distribution of abdominal contents [24], [25]. Physical activity also introduces another source of variability by generating motion artifact and skeletal muscle activity [23], [29]. Together, these effects make posture and physical activity major confounds for ambulatory meal detection systems.

In this study, we introduce a physiology-informed AI-based approach to automated meal detection using HR-EGG and triaxial accelerometry across multiple postural contexts in healthy volunteers as a fully passive, non-invasive readout. We hypothesize that physiologically distinct digestive signatures are sufficiently encoded in HR-EGG and accelerometry to enable automated meal detection even in the context of changing postures. Furthermore, we also hypothesize that posture can affect the signal-to-noise ratio (SNR) of these signatures. Triaxial accelerometry supplies the model with the contextual information needed to disentangle meal-related gastric signatures from noise. By explicitly characterizing how postural transitions modulate gastric myoelectric activity, this study aims to establish a framework for passive, ambulatory dietary monitoring using healthy volunteers as a proof-of-concept.

The remainder of this paper is organized as follows. In Methods, we describe the data collection protocol, signal preprocessing and feature extraction pipeline, and the dilated convolutional neural network architecture. We also detail the feature ablation analysis and the postural analysis. In Results, we report the model’s meal detection performance, the most significant input features from the feature ablation analysis, and the effects of posture on gastric myoelectric signal quality. Finally, in the Discussion, we consider the physiological and clinical implications of the results and outline the study’s limitations and directions for future work.

## Methods

### Dataset

This study was conducted with Institutional Review Board (IRB) approval from the University of California, Berkeley (IRB #2025-11-19143). Seven healthy adult participants (mean age 25.0 years [range 24-28]) completed a total of sixteen standardized meal recording sessions. Relevant demographic information about study participants can be found in Table 1. Participants fasted for a minimum of three hours prior to each session. In seven of the sixteen sessions, the session lasted an average of 150 minutes and consisted of three phases: a 30-minute fasted baseline recording, consumption of a standardized meal, and a 90-minute postprandial monitoring period (Fig. 1). Standardized meals included one of three meals: Barebells cookies and cream protein bar, Trader Joe’s kimbap, or Trader Joe’s handheld pot pie. During the postprandial period, participants completed three 30-minute postural blocks in a randomized order: sitting, walking, and lying down. In the remaining nine sessions, a modified protocol was used in which participants maintained a single fixed posture throughout the entire session rather than all three posture conditions. Together, this allows us to explore the effects of different postures in short and longer spans during digestion.

**Fig 1.**
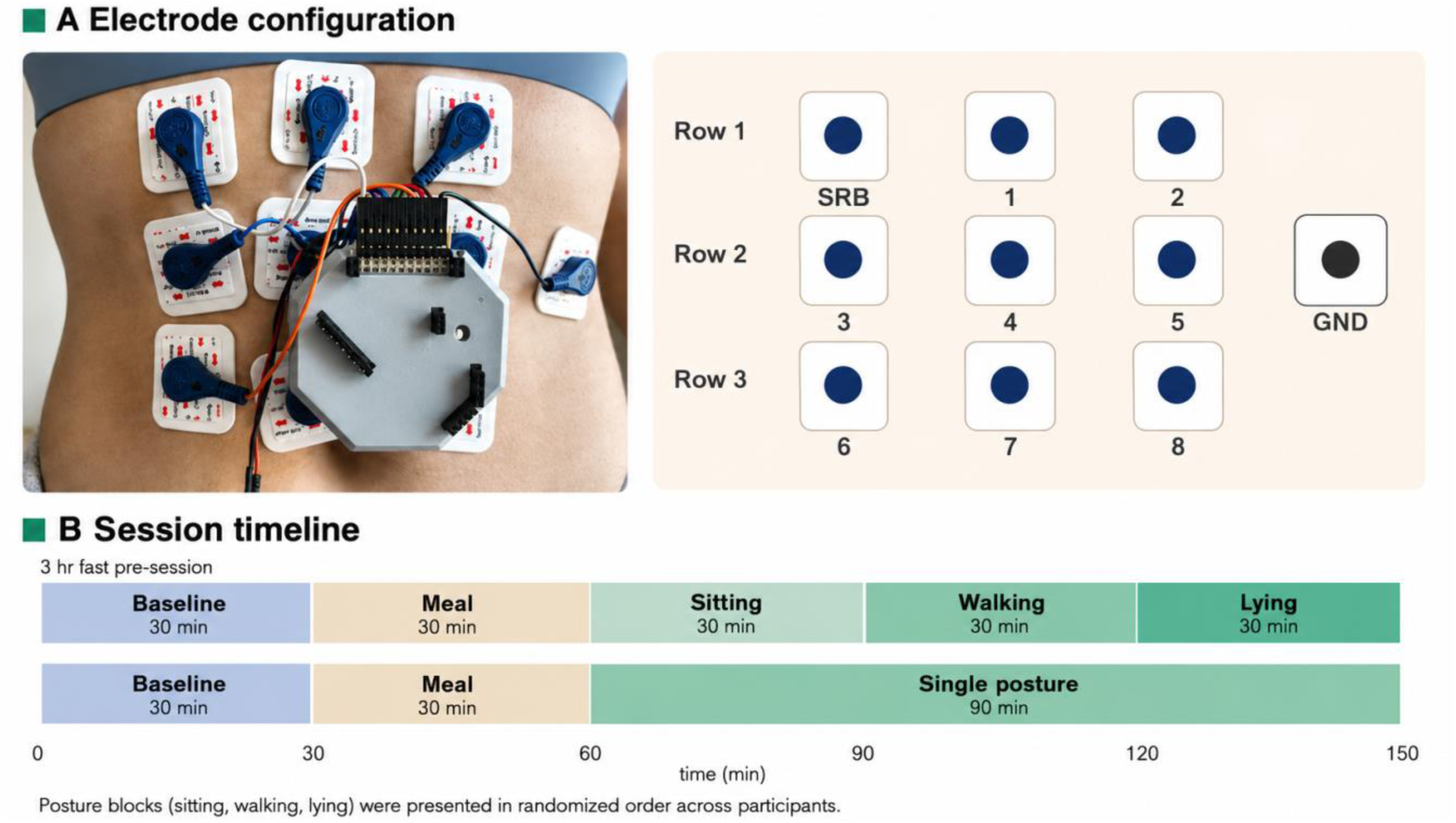
Wearable HR-EGG system electrode placement and session protocol. (A) left: applied electrode array positioned on the abdomen. Right: schematic of the 3×3 electrode grid (channels 1-8 and signal reference bias, SRB) and single ground electrode (GND). Electrodes were arranged in three rows of three over the epigastric region. (B) Each 150-minute recording session consisted of a 30-minute fasted baseline period, consumption of a standardized meal over 30 minutes, and three 30-minute posture blocks (sitting, walking, and lying supine) presented in a randomized order.

**Table 1.** Enrollment characteristics of study participants (N = 7).

| Characteristic | n (%) or Mean [range] |
| --- | --- |
| **Total enrolled** | 7 |
| **Demographics** |  |
| Age, years | 25.0 [24–28] |
| Female sex | 5 (71.4%) |
| **Race / Ethnicity** |  |
| Asian | 5 (71.4%) |
| Two or more races | 2 (28.6%) |
| **Anthropometrics** |  |
| BMI, kg/m<sup>2</sup> | 23.8 [18.3–26.6] |
| **Gastrointestinal Health** |  |
| Self-reported GI score (1–5)<sup>1</sup> | 4.0 [2–5] |

<sup>1</sup> GI score: 1 = daily discomfort, 2 = 2–4 days/week, 3 = once/week, 4 = occasionally, 5 = never or almost never.
BMI = body mass index. Values reported as n (%) or mean [range]. GI = gastrointestinal.

HR-EGG and triaxial accelerometry were recorded simultaneously using an OpenBCI Cyton board [28] at 250 Hz and 25 Hz respectively. Ten Ag-AgCl electrodes, including one reference and one ground electrode, were arranged in a 3×3 spatial grid on the anterior abdominal surface overlying the stomach, with the ground electrode placed on the lateral fatty tissue of the waist (Fig. 1) [29]. Triaxial accelerometry was recorded via the onboard inertial measurement unit (IMU) of the Cyton board, which was physically attached at the electrode positions 4, 5, 7, and 8. We carefully observed and recorded the starting and ending times for the meal.

### Signal Preprocessing and Feature Extraction

Raw HR-EGG signals were decimated from 250 Hz to 1 Hz using a zero-phase finite impulse response (FIR) filter. A per-channel Wiener filter was applied to suppress motion artifacts [30], and all eight Wiener-filtered channels were then averaged to produce a single composite HR-EGG trace (Fig. 2). Similar figures showing all features for each session are shown in the Supplementary Materials (Figs. S1-S16).

**Fig 2.**
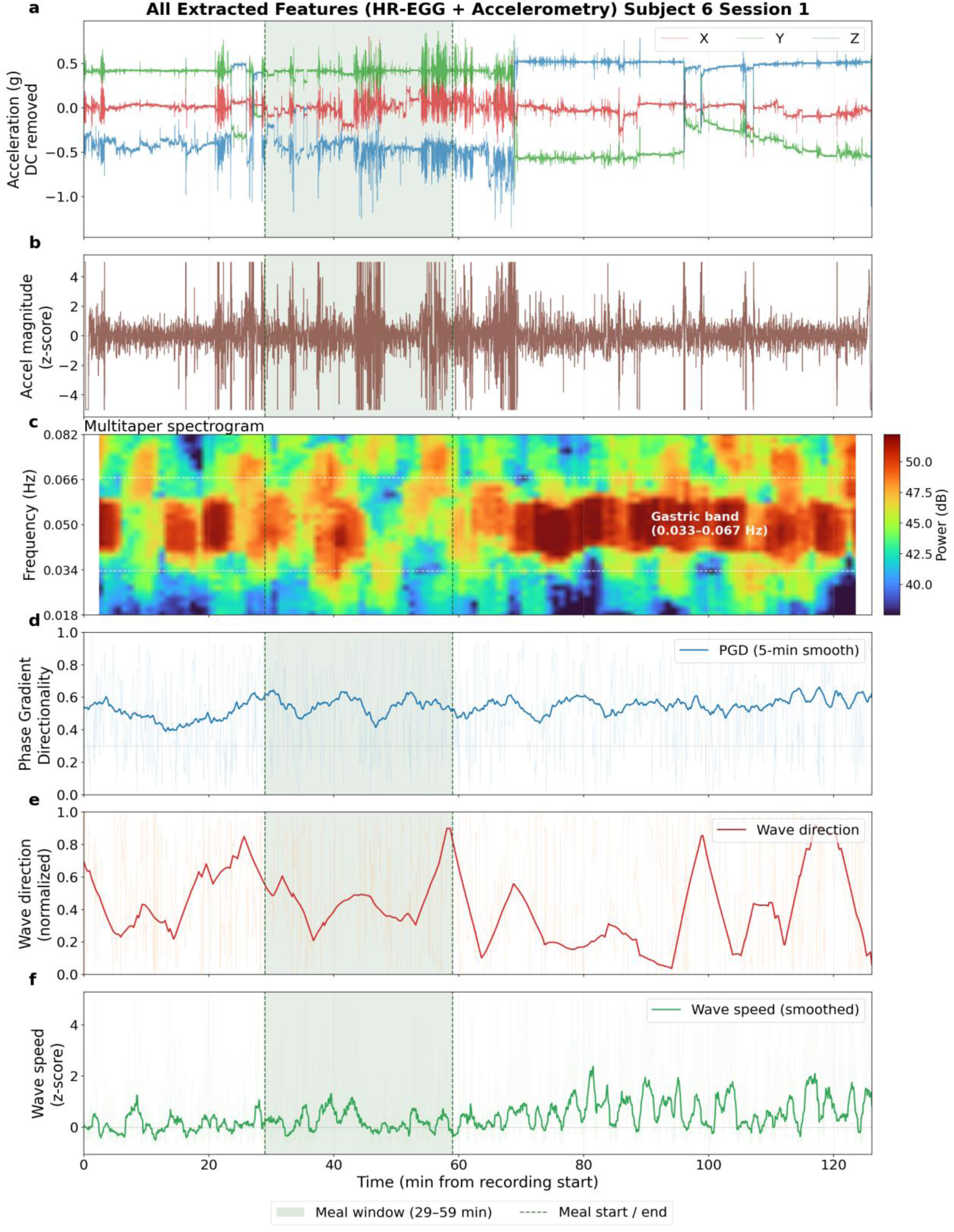
Visualization of all features input into the model from a single representative participant undergoing the three postural changes.. The green window represents the meal period where the participant was actively eating. a) Triaxial accelerometry. (b) Accel magnitude z-score. (c) Multitaper HR-EGG spectrogram. (d) Phase gradient directionality (PGD). (e) Wave direction. (f) Wave speed.

Gastric slow-wave bandpower was estimated using a sliding-window Welch power spectral density (PSD), using a 30-second symmetric Hanning window with 50% overlap. Power was integrated over 2-4 cpm (or equivalently, 0.033-0.067 Hz), corresponding to the physiological range of the normal gastric slow wave [18], [19], [20]. Two representations of the slow-wave were derived from the integrated bandpower: (1) log-transformed absolute bandpower to capture the true postprandial increase in gastric power and (2) session-normalized z-scored bandpower, controlling for inter-subject variability in baseline signal strength.

In addition to the temporal characteristics, three spatial features, phase gradient directionality (PGD), wave propagation direction, and wave propagation speed were computed from the multi-channel HR-EGG data based on previous work [24], [30] after bandpass filtering each channel to the gastric band (2-4cpm) and computing the instantaneous phase using the Hilbert transform [24]. PGD is a value between 0 and 1 that measures the degree of alignment in phase across all channels [24]. Wave speed estimates were set to zero when PGD fell below 0.3 to exclude time points where spatial coherence was insufficient to support reliable propagation estimates (Fig. 2).

Triaxial accelerometry was preprocessed by computing the vector magnitude (2-norm) of acceleration at each time point. A second-order zero-phase Butterworth bandpass filter (0.02-0.45 Hz) was then applied. The 0.02 lower cutoff removed the static gravitational offset and baseline drift associated with postural changes [25], while the upper cutoff of 0.45 was set to be near the Nyquist limit of the 1Hz processed signal. The resulting signal was z-scored within each session.

### Model Architecture

Each session was truncated or zero-padded post-preprocessing to 9,000 samples (150 minutes at 1 Hz) and partitioned into 30 non-overlapping 5-minute windows of 300 samples each. The zero-padded sections were masked and excluded from the loss calculation. A total of six features (five HR-EGG input features of raw and normalized bandpower, PGD, wave direction, wave speed along with accelerometer magnitude) were concatenated across time to form an 1,800-dimensional input vector per window (Fig. 3). A window was assigned a positive meal label if more than 50% of its duration fell within the manually annotated eating window. This yielded 510 total windows across 16 sessions, with an approximate 4-to-1 class imbalance (408 non-meal, 102 meal).

**Fig 3.**
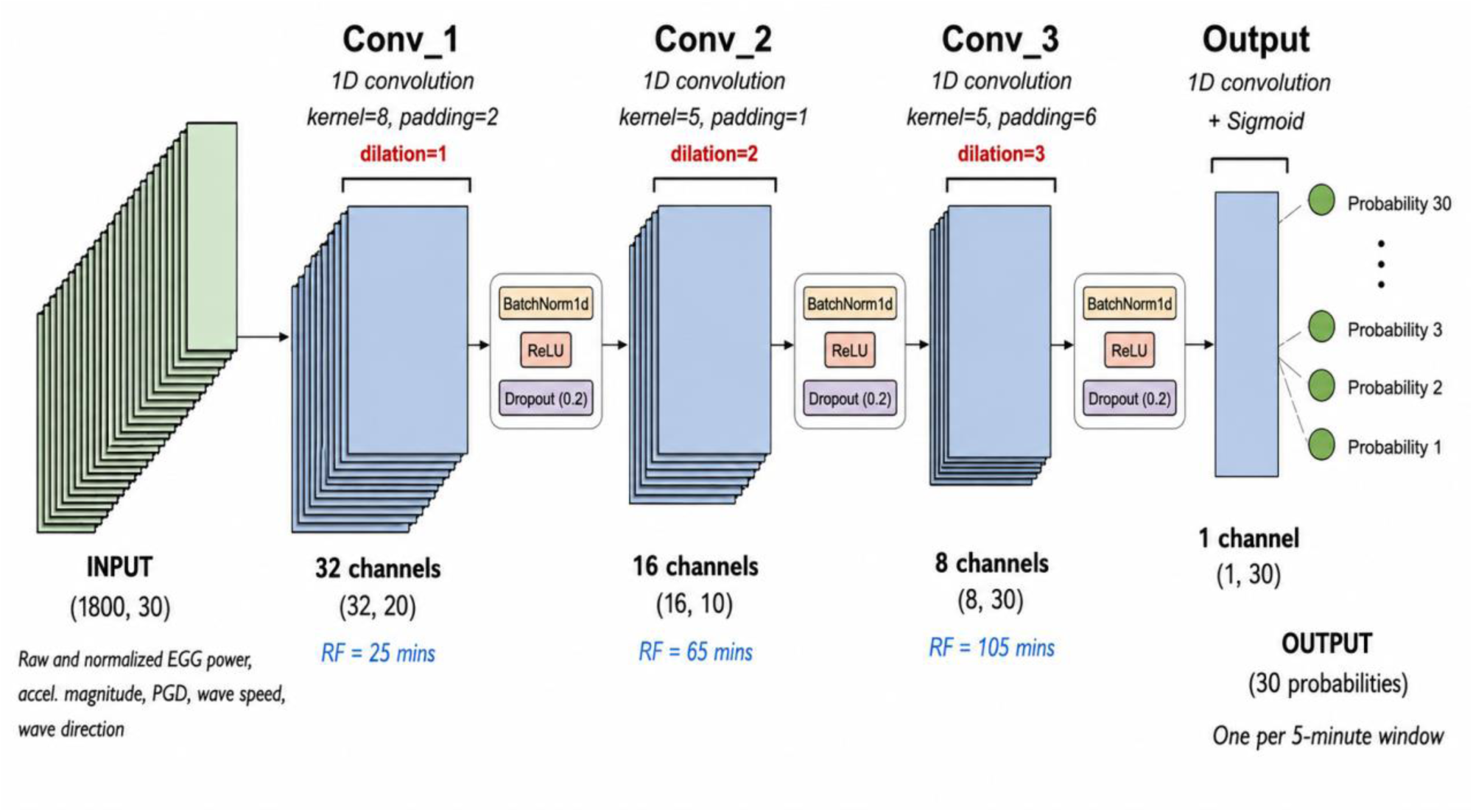
Dilated CNN architecture for automated meal detection using physiological signals. The network processes flatten 5-minute windows of EGG temporal and spatial features and accelerometer magnitude through three progressively dilated convolution blocks (dilation raters: 1, 2, 3) to capture temporal patterns spanning 25 to 105 minutes. Each convolution is followed by normalization, ReLU activation, and dropout regulation. The output layer produces a per-window meal probability across the 150-minute recording.

A one-dimensional dilated convolutional neural network (1D dilated CNN) was trained to classify each window as meal or non-meal (Fig. 3) [31]. The architecture consisted of three successive dilated convolutional blocks followed by a pointwise output layer: Conv1d(1800→32, k=5, dilation=1) → BatchNorm1d → ReLU → Dropout(0.3); Conv1d(32→16, k=5, dilation=2) → BatchNorm1d → ReLU → Dropout(0.3); Conv1d(16→8, k=5, dilation=3) → BatchNorm1d → ReLU → Dropout(0.3); Conv1d(8→1, k=1) → sigmoid. Dilation rates of 1, 2, and 3 were chosen to capture gastric activity across multiple timescales, from the immediate postprandial response through extended digestion.

Model generalization was evaluated using leave-one-subject-out (LOSO) cross-validation across all 7 subjects, training a single model per held-out subject. Performance was then assessed for each of the 16 sessions individually using the model from the corresponding subject’s fold using area under the receiver operating characteristic curve (AUROC), area under the precision-recall curve (AUPRC), and F1-score, with the classification threshold optimized per fold by selecting the value that maximized F1 on the precision-recall curve.

### Feature Importance Analysis

The relative contribution of each input feature to model performance was assessed using a leave-one-feature-out ablation analysis. For each of the six features, all 7 LOSO folds were re-run excluding that single feature while all other hyperparameters remained identical to the primary model. Feature importance was quantified as the change in AUPRC relative to the full model, with a greater decrease in performance indicating greater feature importance.

### Postural Analysis

The 90-minute postprandial period was segmented into three approximately 30-minute postural conditions: sitting, walking, and lying. Nominal phase boundaries were defined as 0, 30, and 60 minutes after the last recorded bite and refined by detecting the sharpest step-change in accelerometer variance within a 5-minute tolerance window.

For each postural condition, four HR-EGG metrics were computed from the best-performing bipolar electrode pair. The best-performing pair was identified by computing the pairwise bipolar derivations (channel differences) for all 28 channel pairs and selecting the pair with the highest normalized power in the gastric band (0.033-0.067 Hz). Spectral features were derived from a 4-minute multitaper spectrogram with 75% overlap and a 1-minute step. **Spectral signal-to-noise ratio** (SNR, in dB) was defined as the power in the gastric band versus the power in the band immediately above (0.07-0.2 Hz) as a measure of noise. **Dominant frequency** (cpm) was taken as the mode of the peak frequency across spectrogram windows within each condition. The **percentage of normogastric activity** was defined as the fraction of windows with a dominant peak within the normogastric range (0.033-0.067 Hz). Mean PGD was computed as a measure of spatial wavefront coherence per condition. Signal envelope stability within each postural condition was characterized by **the number of sustained waves**, which was defined as the count of continuous intervals in which PGD exceeded 0.5 for at least 60 consecutive seconds within each condition [24]. The gastric band amplitude envelope was computed by bandpass filtering each channel to the 0.033-0.067 Hz and extracting the Hilbert amplitude. Differences across the three postural conditions were assessed using the Friedman test, a non-parametric repeated-measures test to account for the small sample size and repeated subject testing [32].

## Results

### Model Performance

Table 2 summarizes the performance metrics and session characteristics for all 16 sessions across seven participants. Fig 4 displays the two best-performing sessions and the two worst-performing sessions. Table 3 presents the results of the leave-one-feature-out ablation analysis, ranked by change in AUPRC. Table 4 reports the gastric myoelectric metrics by postural conditions, and Fig 5 provides a representative visualization of the HR-EGG bandpower for each posture.

**Fig 4.**
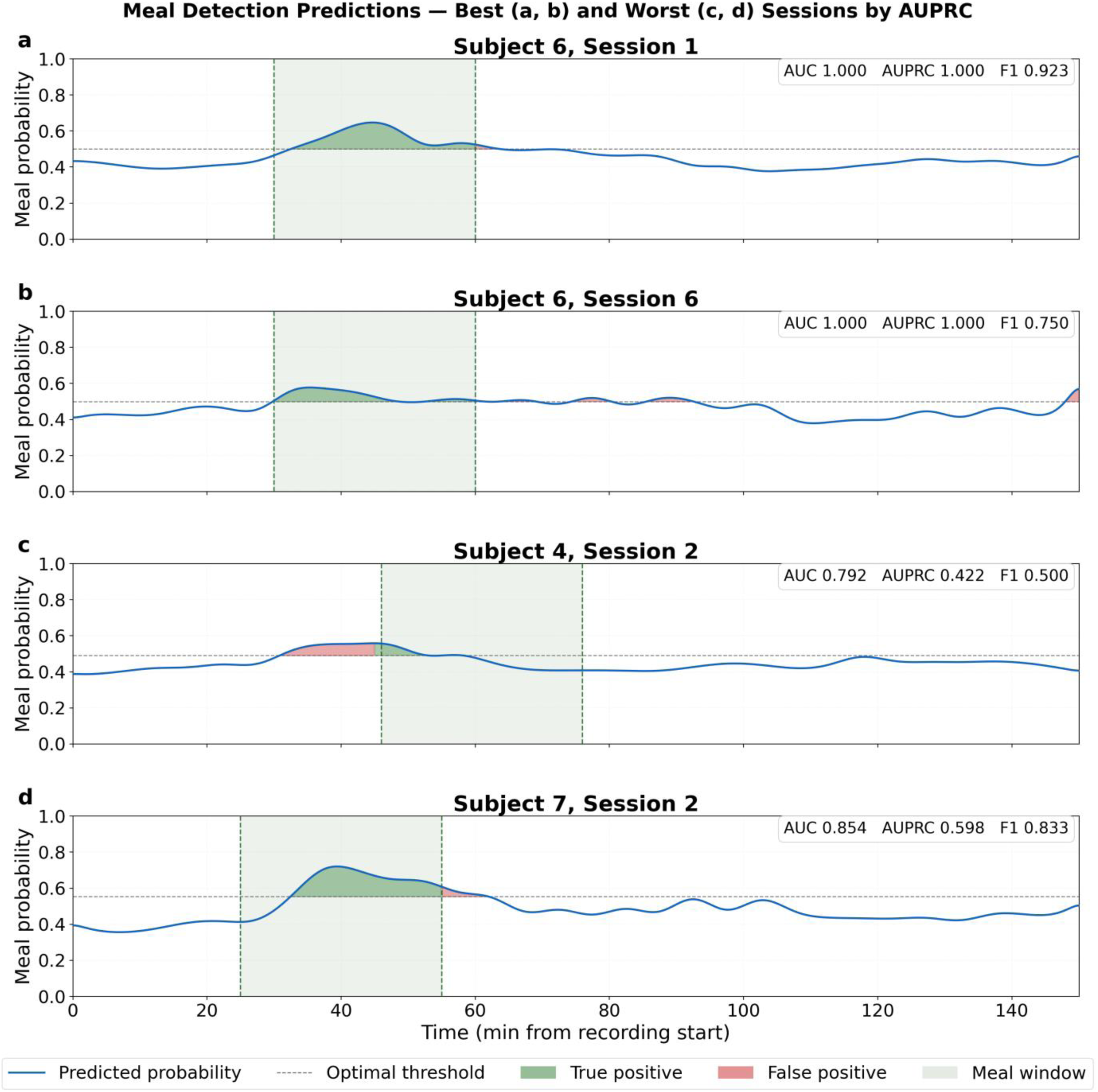
Smoothed meal probability outputs from the dilated 1D CNN are shown for the two highest-performing sessions by AUPRC (panels a, b) and the two lowest-performing sessions (panel c, d). Each panel represents one held-out test subject under leave-one-subject-out cross-validation, with the model trained on the remaining six sessions. The blue curve shows the predicted meal probability; the dashed horizontal line indicates the optimal classification threshold for each session, determined by maximizing F1 scores on the precision-recall curve. Green shading indicates true positive regions, whereas red shading indicates false positive regions. The light green shaded region between the dashed vertical boundaries marks the manually annotated consumption window.

**Fig 5.**
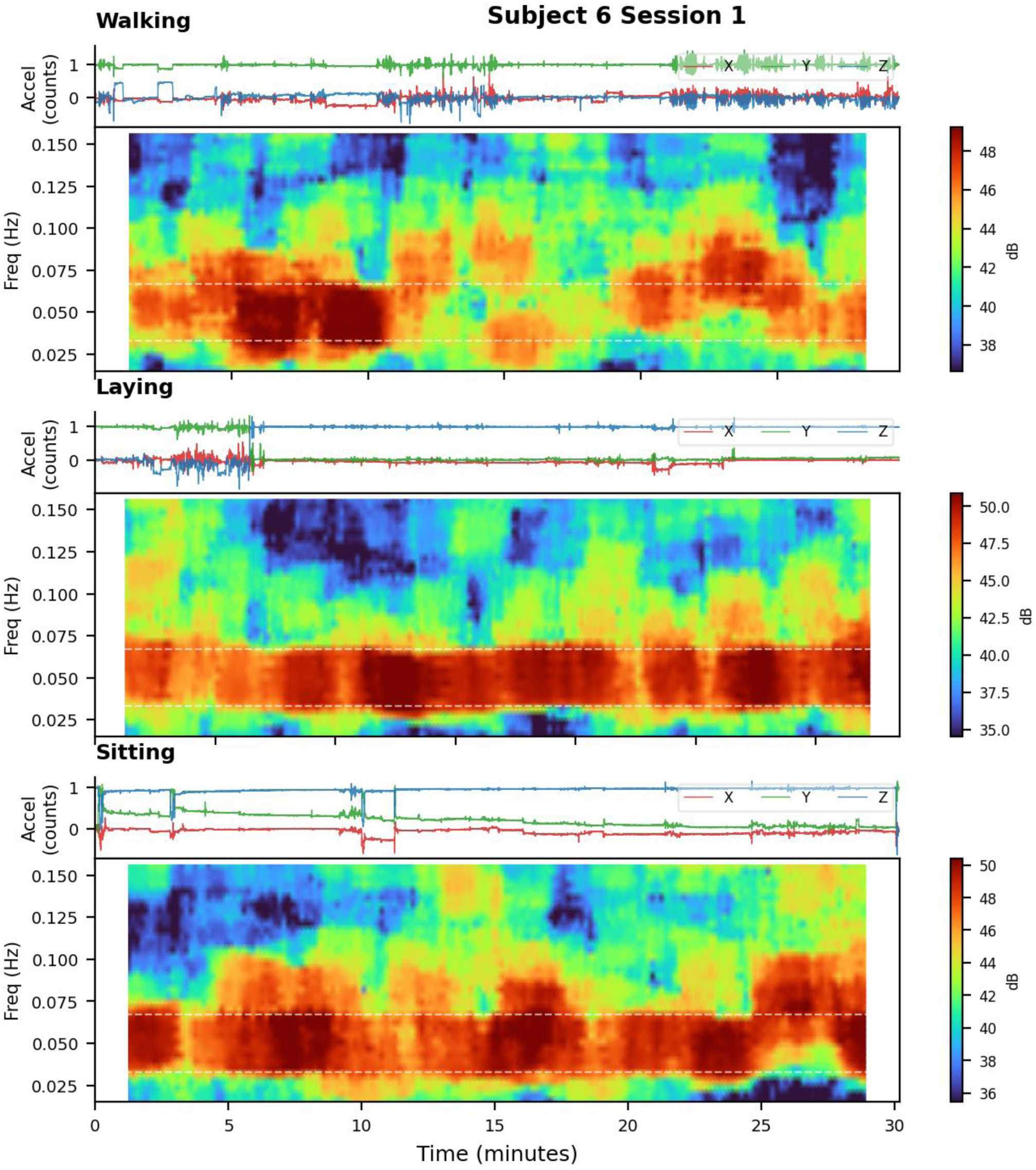
Representative postprandial recordings across three postural conditions. Each condition panel shows triaxial accelerometry (top: X, Y, Z) and a single-channel multitaper HR-EGG spectrogram of bandpower (bottom) from the corresponding post-meal phase.

**TABLE 2.** Per-Subject Classification Performance, Sorted by AUROC.

| Subject | AUROC | AUPRC | F1 | Threshold | Meal | Session (mins) | Meal (mins) | Posture Order |
| --- | --- | --- | --- | --- | --- | --- | --- | --- |
| S6 | 1.000 | 1.000 | 0.909 | 0.556 | 600 kcal | 150 | 50 | Sit, Walk, Lie |
| S6 | 1.000 | 1.000 | 0.909 | 0.498 | 200 kcal | 165 | 29 | Lie, Walk, Sit |
| S3 | 0.993 | 0.976 | 0.824 | 0.493 | 600 kcal | 129 | 16 | Walk, Lie, Sit |
| S1 | 0.972 | 0.933 | 0.800 | 0.681 | 400 kcal | 122 | 30 | Sit, Lie, Walk |
| S6 | 0.972 | 0.911 | 0.769 | 0.402 | 400 kcal | 196 | 29 | Sit, Lie, Walk |
| S6 | 0.972 | 0.897 | 0.769 | 0.460 | 200 kcal | 139 | 28 | Sit, Lie, Walk |
| S6 | 0.965 | 0.906 | 0.727 | 0.402 | 600 kcal | 154 | 19 | Sit, Walk, Lie |
| S6 | 0.958 | 0.863 | 0.714 | 0.531 | 600 kcal | 192 | 35 | Lie, Walk, Sit |
| S6 | 0.944 | 0.862 | 0.667 | 0.641 | 600 kcal | 126 | 28 | Sit, Walk, Lie |
| S7 | 0.938 | 0.900 | 0.800 | 0.454 | 200 kcal | 151 | 27 | Sit, Lie, Walk |
| S5 | 0.917 | 0.641 | 0.667 | 0.456 | 600 kcal | 120 | 30 | Sit, Lie, Walk |
| S5 | 0.896 | 0.881 | 0.800 | 0.549 | 200 kcal | 160 | 30 | Stand, Lie, Sit |
| S7 | 0.854 | 0.598 | 0.600 | 0.535 | 200 kcal | 157 | 58 | Lie, Sit, Walk |
| S4 | 0.824 | 0.698 | 0.500 | 0.530 | 400 kcal | 138 | 30 | Walk, Sit, Lie |
| S4 | 0.792 | 0.422 | 0.526 | 0.512 | 600 kcal | 156 | 30 | Sit, Stand, Lie |
| S2 | 0.799 | 0.693 | 0.600 | 0.629 | 600 kcal | 112 | 15 | Lie, Sit, Walk |
| Average | 0.925 | 0.824 | 0.724 | 0.520 | - | 147.94 | 30.18 | - |

AUROC = area under the receiver operating characteristic curve; AUPRC = area under the precision-recall curve; F1 = F1 score at optimal threshold. A null (chance) classifier yields AUROC = 0.5 for all sessions by definition; AUPRC null equals the meal-window prevalence (mean 0.20 across all sessions). Meal caloric content: protein bar = 200 kcal, kimbab = 400 kcal, potpie = 600 kcal. Posture order reflects randomized sequence during the session.

**TABLE 3.** Leave-One-Out Feature Ablation Analysis.

| Feature Removed | AUPRC<br>Mean null:<br>0.20 | $\Delta\text{AUPRC}$ | AUROC<br>null: 0.50 | $\Delta\text{AUROC}$ | F1-Score | $\Delta\text{F1}$ |
| --- | --- | --- | --- | --- | --- | --- |
| **Full model (6 features)** | **0.824** | — | **0.925** | — | **0.724** | — |
| Phase gradient directionality (PGD) | 0.636 | -0.188 | 0.792 | -0.133 | 0.706 | -0.018 |
| Normalized EGG bandpower | 0.650 | -0.174 | 0.824 | -0.101 | 0.737 | +0.013 |
| Accelerometer magnitude | 0.655 | -0.169 | 0.808 | -0.117 | 0.709 | -0.015 |
| Wave propagation direction | 0.661 | -0.163 | 0.822 | -0.103 | 0.719 | -0.005 |
| Raw EGG bandpower | 0.692 | -0.132 | 0.827 | -0.098 | 0.744 | +0.020 |
| Wave propagation speed | 0.719 | -0.105 | 0.862 | -0.063 | 0.741 | +0.017 |

$\Delta$ values are relative to the full six-feature model within the same leave-one-subject-out cross-validation run. A null model would yield an AUROC of 0.50 and an AUPRC of 0.20, where 0.20 reflects the average positive class proportion in the dataset. Rows sorted by $\Delta\text{AUPRC}$ .

**TABLE 4.**
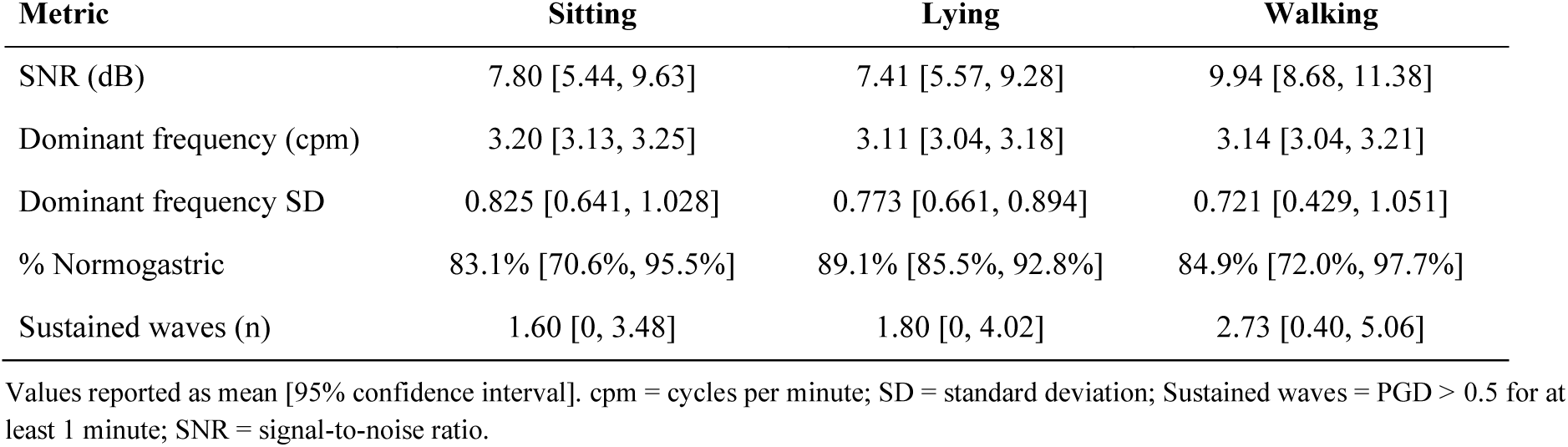
Gastric Myoelectric Metrics by Postural Condition (Mean, 95% CI)

Model performance was evaluated using leave-one-subject-out cross-validation across the seven subjects, with performance reported for each of the 16 sessions individually (Table 2).

The full six-feature model achieved a mean AUROC of 0.925 ± 0.073 (null model = 0.50), mean AUPRC of 0.824 ± 0.169 (mean null model = 0.20) and mean F1 score of 0.724 ± 0.131 across the 16 sessions (Table 2). Individual session AUROC ranged from 0.799 to 1.000, with three sessions achieving perfect or near-perfect discrimination (AUROC ≥ 0.993) (Fig. 4). Similar performance figures for all session can be found in Supplementary Materials (Figs. S17-S32). AUPRC ranged from 0.422 to 1.00, and F1 from 0.500 to 0.909 across all sessions. The weakest performance was concentrated in subject 4, whose two sessions yielded among the lowest AUROC (0.792 and 0.824) and AUPRC (0.422 and 0.698) values, with one additional low-performing session from subject 2 (Fig. 4). No consistent relationship was observed between meal size (200–600 kcal) and classification performance.

### Feature importance

Feature importance was assessed primarily through the change in AUPRC, given the substantial class imbalance in the dataset (Table 3). Removal of phase gradient directionality (PGD) produced the largest decrease in AUPRC relative to the full model (ΔAUPRC = −0.188; ΔAUROC = −0.133, ΔF1 = −0.018), identifying it as the single most informative feature. Removal of normalized EGG bandpower produced the second-largest AUPRC decrease (ΔAUPRC = −0.174), followed by accelerometer magnitude (ΔAUPRC = −0.169) and wave propagation direction (ΔAUPRC = −0.163). Removal of raw EGG bandpower produced a smaller decrease (ΔAUPRC = −0.132), and removal of wave propagation speed produced the smallest decrease (ΔAUPRC = −0.105, ΔAUROC = −0.063). For every feature, removal reduced both AUPRC and AUROC, indicating that all six inputs contributed positively to classification, with PGD contributing most and wave propagation speed least. Across features, AUPRC decreases were consistently larger in magnitude than AUROC decreases.

### Postural effects on gastric myoelectric activity

Posture order varied across sessions and subjects (Table 4). Accelerometer RMS confirmed that the activity manipulation was successful, with walking producing the highest values (0.412 g), followed by sitting (0.346 g) and lying (0.253 g) (Fig. 5). Postural figures for all sessions can be found in the Supplementary Materials (Figs. S33-S39).

Mean spectral SNR was highest during walking (9.94 dB [95% CI: 8.68, 11.38]) compared to sitting (7.80 dB [5.44, 9.63]) and lying (7.41 dB [5.57, 9.28]) (Table 4). The percentage of normogastric activity (2–4 cpm) was high across all conditions, with lying showing the greatest consistency (89.1% [85.5%, 92.8%]), followed by walking (84.9% [72.0%, 97.7%]) and sitting (83.1% [70.6%, 95.5%]). Dominant frequency remained within the normogastric range across all three conditions (sitting: 3.20 cpm [3.13, 3.25]; lying: 3.11 cpm [3.04, 3.18]; walking: 3.14 cpm [3.04, 3.21]), with no meaningful difference across conditions. Spatial wavefront coherence was indexed by the number of sustained waves, defined as continuous intervals in which PGD exceeded 0.5 for at least 60 seconds in the full 30 minutes of each position. The mean number of sustained waves per session ranged from 1.6 (sitting, 95% CI [0, 3.5]) to 2.7 (walking, 95% CI [0.4, 5.1]), with considerable inter-subject variability since individual session counts ranged from 0 to 6. Friedman tests across the three within-session postural conditions (n = 7 sessions with complete postural data) did not reach statistical significance for any metric (SNR: p = 0.368; dominant frequency: p = 0.092; % normogastric: p = 1.000; sustained waves: p = 0.405).

## Discussion

In this study, we developed a physiology-informed machine learning framework for automated meal detection using HR-EGG and triaxial accelerometry data from seven healthy volunteers across 16 standardized meal sessions including multiple postural contexts. We found that our model, a dilated one-dimensional convolutional neural network, detected active meal consumption using six primary features at five-minute resolution with mean AUROC of 0.925 ± 0.073 and mean AUPRC of 0.824 ± 0.169, evaluated using leave-one-subject-out cross-validation. Furthermore, using a feature ablation analysis, we found that PGD was the most important single feature and wave speed was the least. In parallel, we characterized how posture modulates gastric myoelectric signal quality to assess the feasibility of meal detection under the ambulatory conditions that have historically limited electrogastrography. Across the three postural conditions, walking produced the highest SNR while lying showed the most consistent normogastric rhythm, and dominant frequency remained stable regardless of posture.

There are three key takeaways from this work. First, our full model performance (AUROC of 0.925 ± 0.073, AUPRC of 0.824 ± 0.169) supports our hypothesis that HR-EGG data and accelerometry can be used for automated meal detection. While there was variability in performance across meal size, posture order, and recording conditions, the AUPRC alone, highly relevant for an imbalanced dataset, represents a roughly fourfold improvement over the naïve classifier (null model AUPRC of ∼0.20). These results are especially notable given the difficulty of the classification task compared to the minimum to be useful. For example, a coarse binary output determining whether a meal occurred at any point in a large window of time, such as a 30-minute increment, would by itself constitute the basic definition of ‘automated meal detection’, especially for purposes of reducing human recall bias and error. However, we trained the model to identify the precise temporal boundaries of active meal consumption at a 5-minute resolution and penalized for detections that were mistimed, incomplete, or preceded actual ingestion. This is a significantly more challenging task than the bare minimum. The mean F1 of 0.724 ± 0.131 should also be interpreted in that context, as it reflects a stricter performance criterion than would be required for a basic meal detection model. This temporal precision suggests that there is more information that can be gleaned from HR-EGG and accelerometry, such as meal pacing or even estimation of size of meal from duration.

Second, the feature ablation analysis identified PGD as the single most informative feature (ΔAUPRC = −0.188, ΔAUROC = −0.133), with its removal producing the largest decrease in AUPRC, which is the most relevant metric in this imbalanced dataset. PGD as the primary feature is consistent with the broader case for HR-EGG over traditional single-channel EGG, since spatial information can only be captured with multiple channels and improves upon purely temporal feature-based models [24]. Removal of normalized EGG bandpower, accelerometer magnitude, and wave propagation direction produced comparable AUPRC decreases (ΔAUPRC = −0.174, −0.169, and −0.163, respectively), indicating that each contributes meaningfully to detection. The contribution of HR-EGG bandpower is consistent with prior HR-EGG literature [19], [20], [21]. Across all six features, removal reduced both AUPRC and AUROC, indicating that every input contributed positively to classification. Wave propagation speed was the least informative, with its removal producing the smallest decreases (ΔAUPRC = −0.105, ΔAUROC = −0.063) and a marginal increase in F1 (ΔF1 = +0.017). This suggests that while speed carries some discriminative information, the computation of wave propagation speed may be disproportionately susceptible to noise under ambulatory conditions.

Third, posture modulated gastric signal quality in physiologically interpretable ways. Walking produced the highest spectral SNR (9.94 dB) and sustained waves (2.73) relative to sitting (7.80 dB) and lying (7.41 dB), a pattern that was directionally consistent across sessions despite not reaching statistical significance at this sample size. This finding is consistent with prior literature demonstrating that light to moderate physical activity accelerates gastric emptying and enhances gastric motility relative to sedentary activities, possibly through exercise-associated gastrointestinal peptide release and enteric nervous system activation [33], [35]. In contrast, lying down showed the lowest SNR but the highest percentage of normogastric activity (89.1%), suggesting that the supine posture keeps the slow wave propagating at a consistent, regular frequency, which is governed by the ICC pacemaker network and is relatively insensitive to autonomic state [18], [20], [24]. The absence of movement and the extended, uncompressed position of the torso when lying supine may also reduce motion artifact and improve electrode contact. Sitting showed the lowest stability across most metrics, including the fewest sustained waves, which may reflect the tendency of seated subjects to adopt a slouched posture that compresses the torso and degrades electrode contact and signal quality. Across all three conditions, dominant frequency remained within the normogastric range (3.11-3.20 cpm), confirming that postural context modulates gastric signal amplitude and organization rather than intrinsic slow-wave frequency.

This study has several limitations. First, the cohort was small and demographically narrow with seven young adults (mean age 25.0 years, predominantly female), which limits the diversity of physiological responses to which the model was exposed. Larger and more heterogeneous cohorts in future work will improve generalizability. Additionally, this study was a proof-of-concept in healthy volunteers, whereas the clinical conditions of greatest interest, diabetes, gastroparesis, and functional dyspepsia, all involve altered gastric myoelectric activity, so the framework’s performance in clinical populations remains to be established. Validating the approach in these patient groups and in larger, more diverse cohorts is the key next step towards real-world translation, where a passive, non-invasive readout of postprandial gastric activity can surpass existing diagnostic and monitoring tools for automated meal detection.

## Conclusion

This study demonstrates that automated meal detection is achievable in ambulatory settings using a physiology-informed dilated 1D CNN applied to HR-EGG and triaxial accelerometry, achieving a mean AUROC of 0.925 and mean AUPRC of 0.824 across sixteen meals and seven healthy volunteers in a multi-posture recording. By treating gastric myoelectric activity as a continuous digital biomarker of digestive state rather than a static measurement, this framework takes a meaningful step toward passive, objective dietary monitoring that does not depend on manual logging.

## List of abbreviations

HR-EGG: high-resolution electrogastrography
CGM: continuous glucose monitor
GES: gastric emptying scintigraphy
ICC: interstitial cells of Cajal
cpm: cycles per minute
EGG: electrogastrography
SNR: signal-to-noise ratio
IRB: institutional review board
IMU: inertial measurement unit
PSD: power spectral density
FIR: finite impulse response
PGD: phase gradient directionality
CNN: convolutional neural network
LOSO: leave-one-subject-out
AUROC: area under the receiver operating characteristic curve
AUPRC: area under the precision-recall curve
RMS: root mean square
CI: confidence interval
BMI: body mass index
SD: standard deviation

## Declarations

## Ethics approval and consent to participate

This study was conducted with Institutional Review Board (IRB) approval from the University of California, Berkeley (IRB #2025-11-19143). All participants provided written informed consent in accordance with the IRB protocol.

## Consent for publication

The manuscript does not include any individual person’s identifiable data in any form. All participants provided written informed consent in accordance with the IRB protocol.

## Availability of data and materials

The data will be in the PhysioNet repository and code will be in GitHub. Both will be made publicly available upon publication.

## Authors’ contributions

B.C. was involved in conceptualization, data curation, formal analysis, investigation, methodology, software, validation, visualization, writing the original draft and revision and editing of the manuscript. S.S. was involved in conceptualization, funding acquisition, investigation, supervision, and revision and editing of the manuscript.

## Acknowledgements

B.C. and S.S. would like to acknowledge Dr. Connie Wang and Christopher Seaman for their feedback and guidance.

## Endnotes

There are no endnotes.

