## Supplementary Materials for "“Towards Passive Dietary Monitoring: Dilated CNN-Based Meal Detection Using Ambulatory High-Resolution Electrogastrography”"

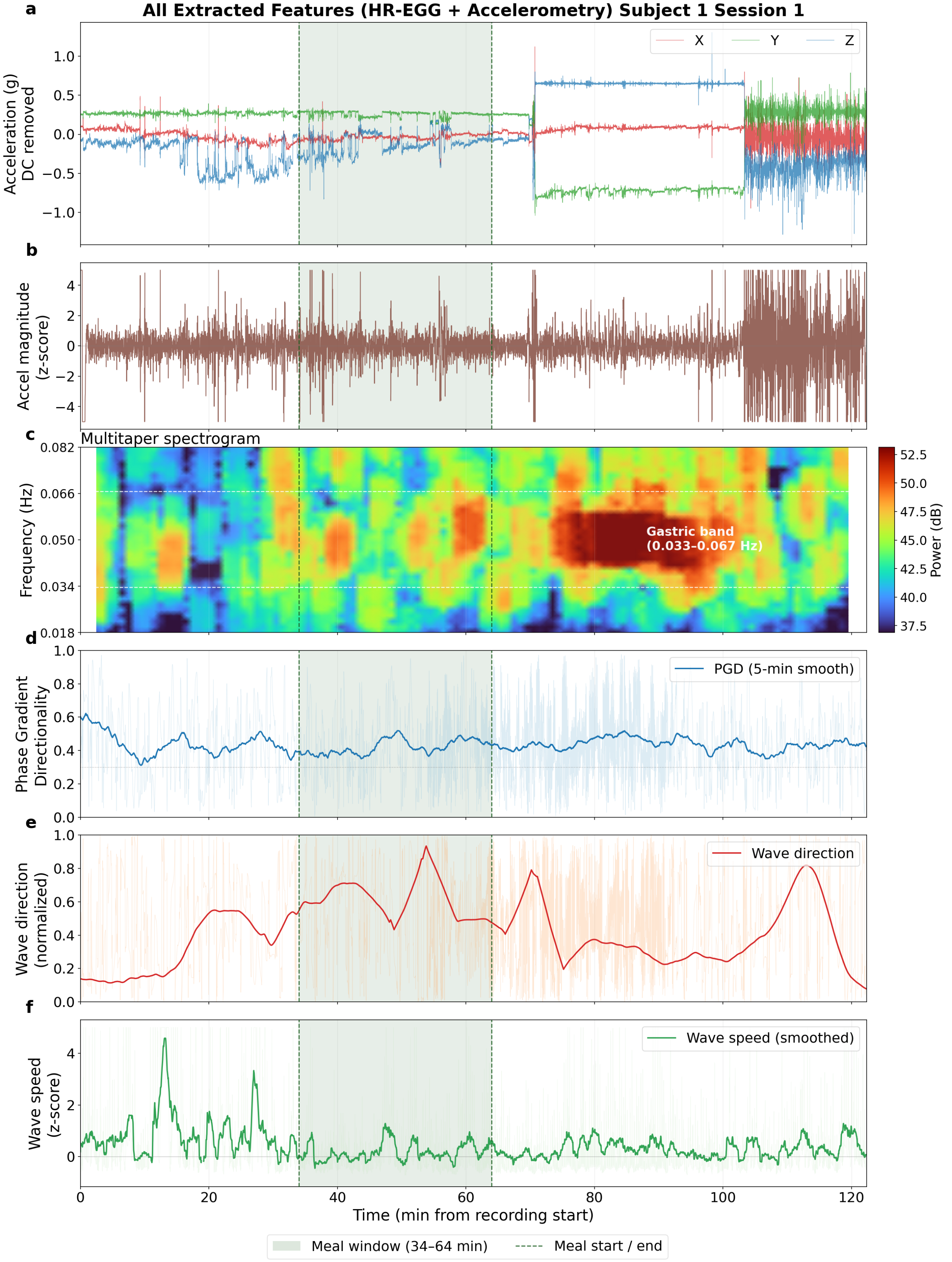


Figure S1. (Subject S1). (a) Triaxial accelerometry. (b) Accel magnitude z-score. (c) Multitaper HR-EGG spectrogram. (d) Phase gradient directionality (PGD). (e) Wave direction. (f) Wave speed.

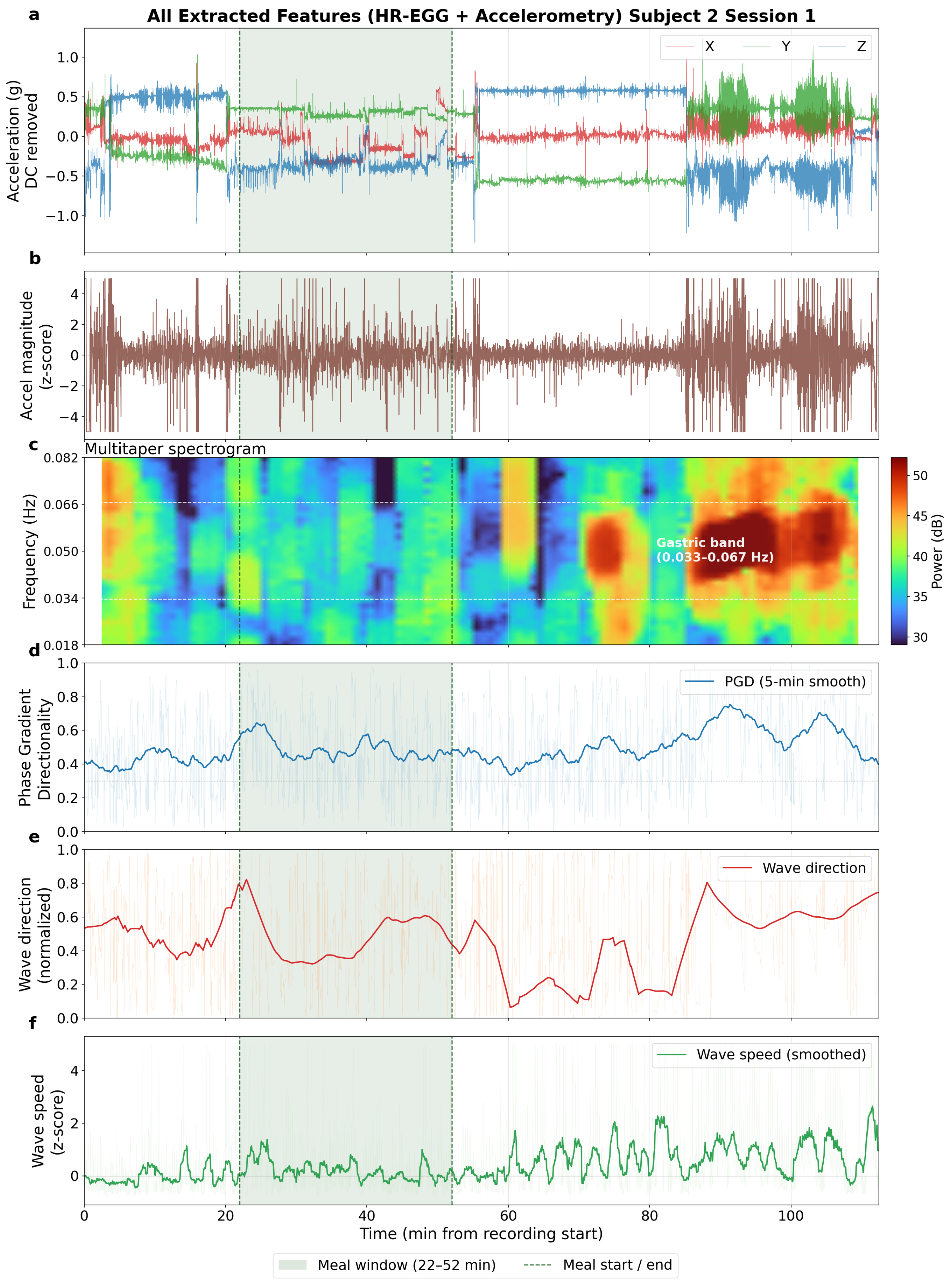


Figure S2. (Subject S2). (a) Triaxial accelerometry. (b) Accel magnitude z-score. (c) Multitaper HR-EGG spectrogram. (d) Phase gradient directionality (PGD). (e) Wave direction. (f) Wave speed.

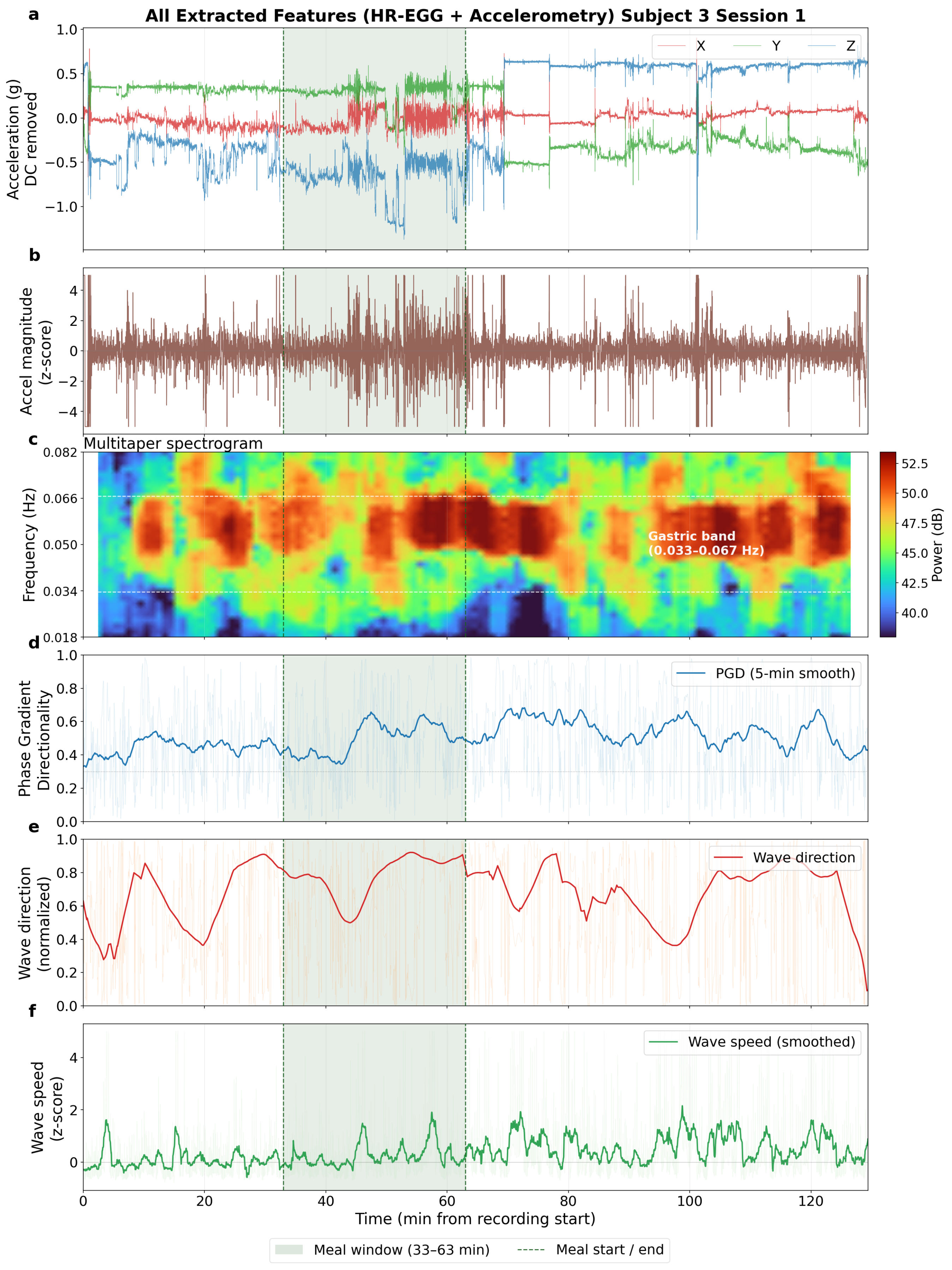


Figure S3. (Subject S3). (a) Triaxial accelerometry. (b) Accel magnitude z-score. (c) Multitaper HR-EGG spectrogram. (d) Phase gradient directionality (PGD). (e) Wave direction. (f) Wave speed.

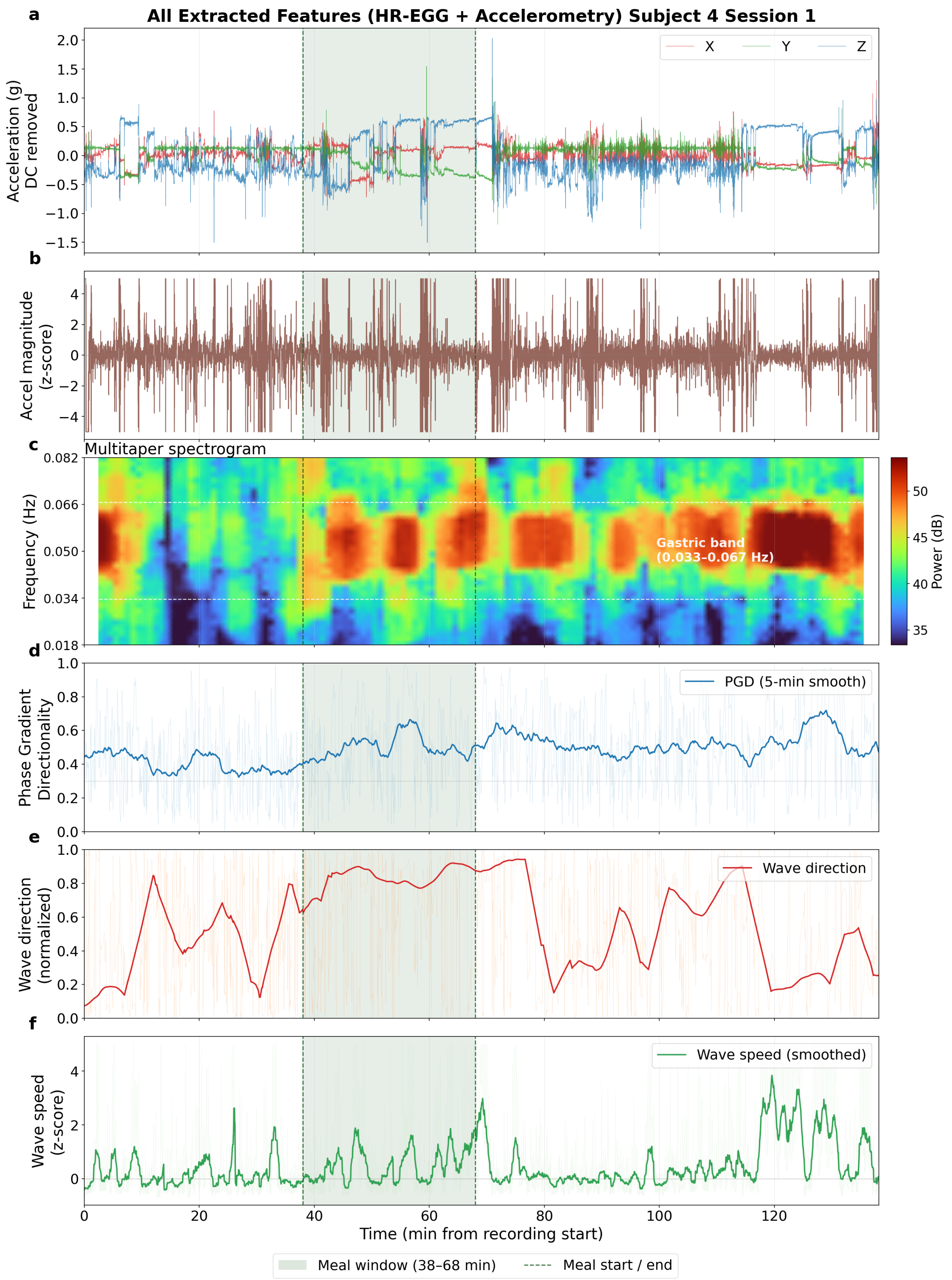


Figure S4. (Subject S4). (a) Triaxial accelerometry. (b) Accel magnitude z-score. (c) Multitaper HR-EGG spectrogram. (d) Phase gradient directionality (PGD). (e) Wave direction. (f) Wave speed.

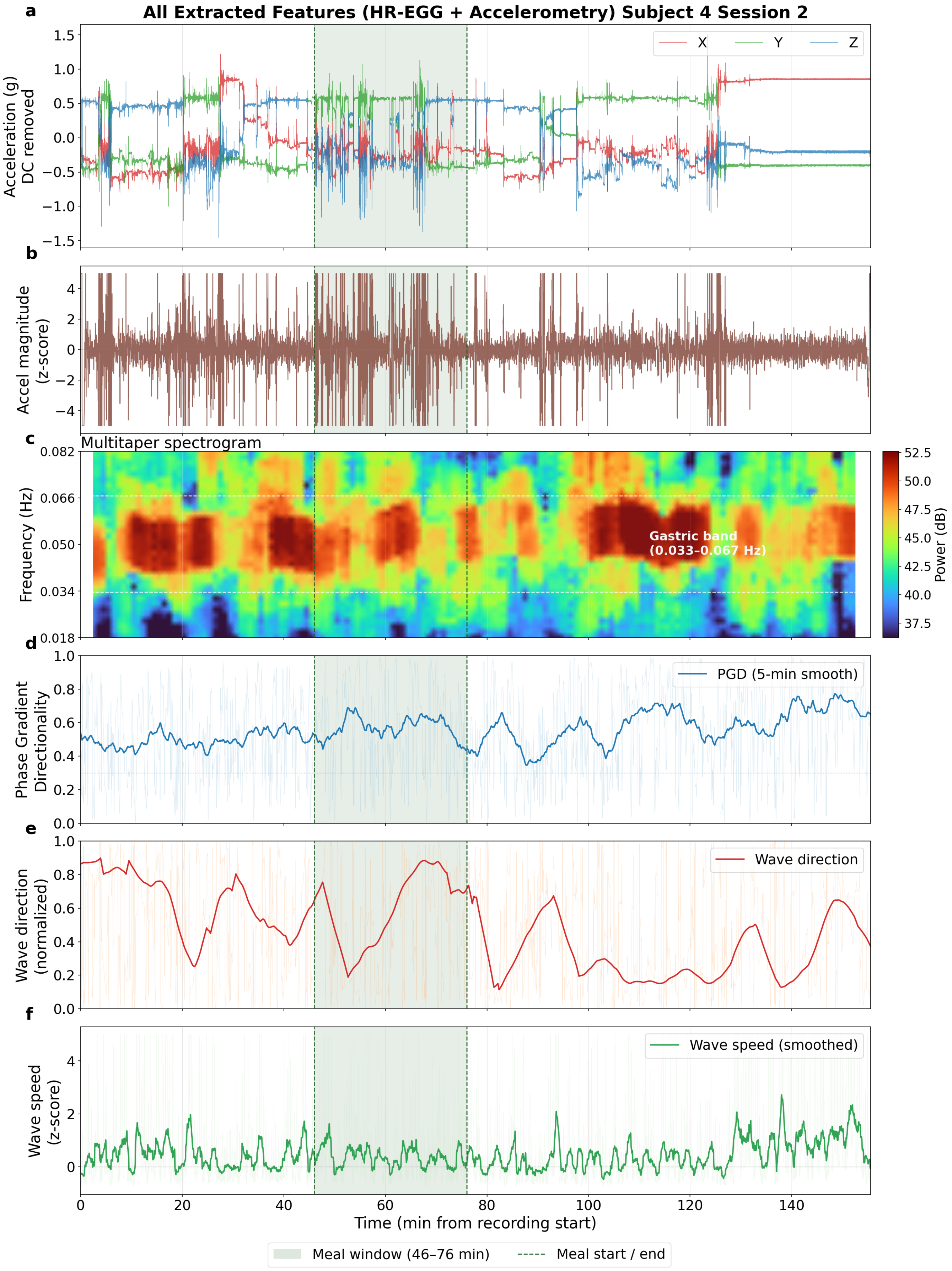


Figure S5. (Subject S4). (a) Triaxial accelerometry. (b) Accel magnitude z-score. (c) Multitaper HR-EGG spectrogram. (d) Phase gradient directionality (PGD). (e) Wave direction. (f) Wave speed.

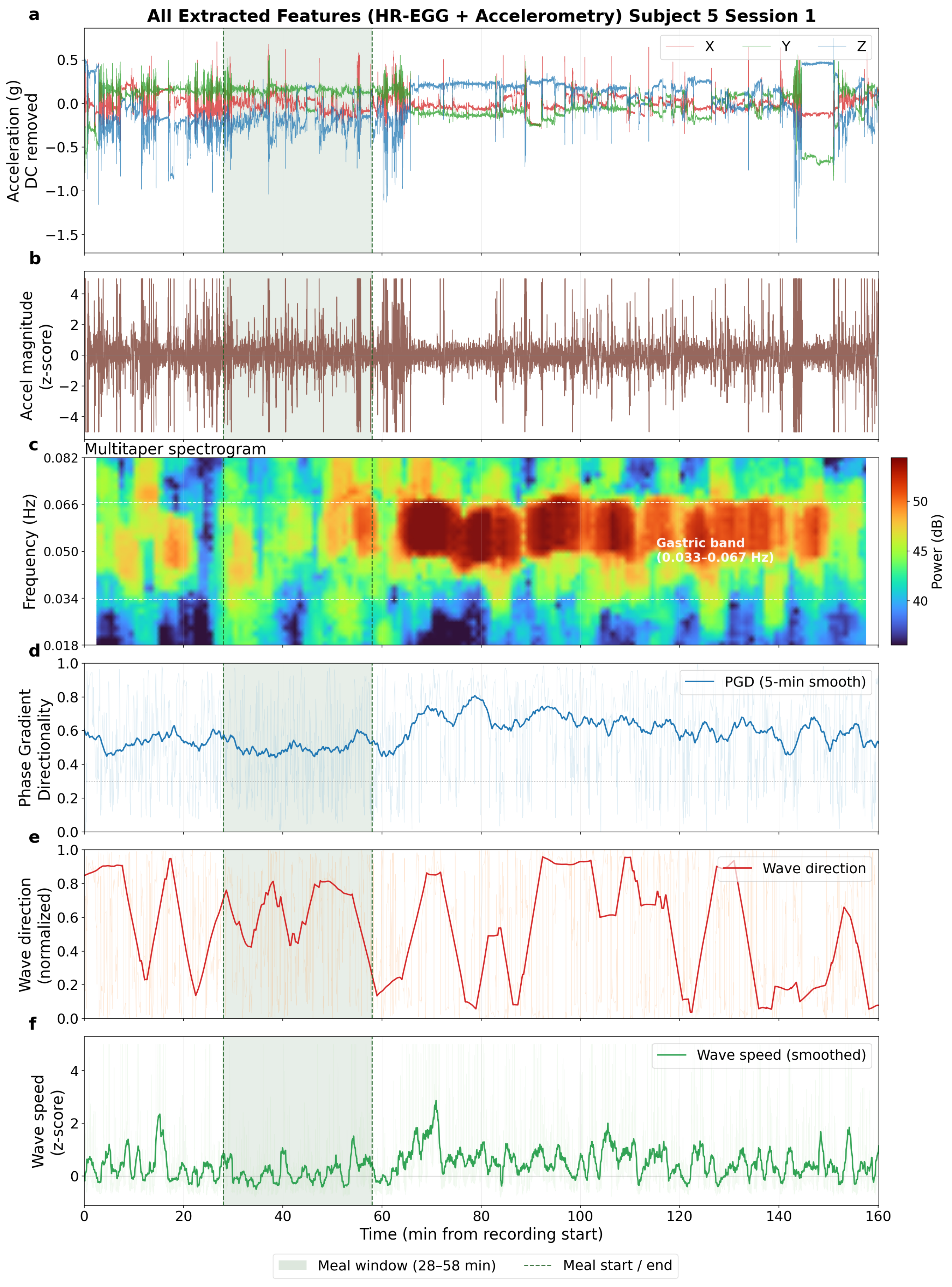


Figure S6. (Subject S5). (a) Triaxial accelerometry. (b) Accel magnitude z-score. (c) Multitaper HR-EGG spectrogram. (d) Phase gradient directionality (PGD). (e) Wave direction. (f) Wave speed.

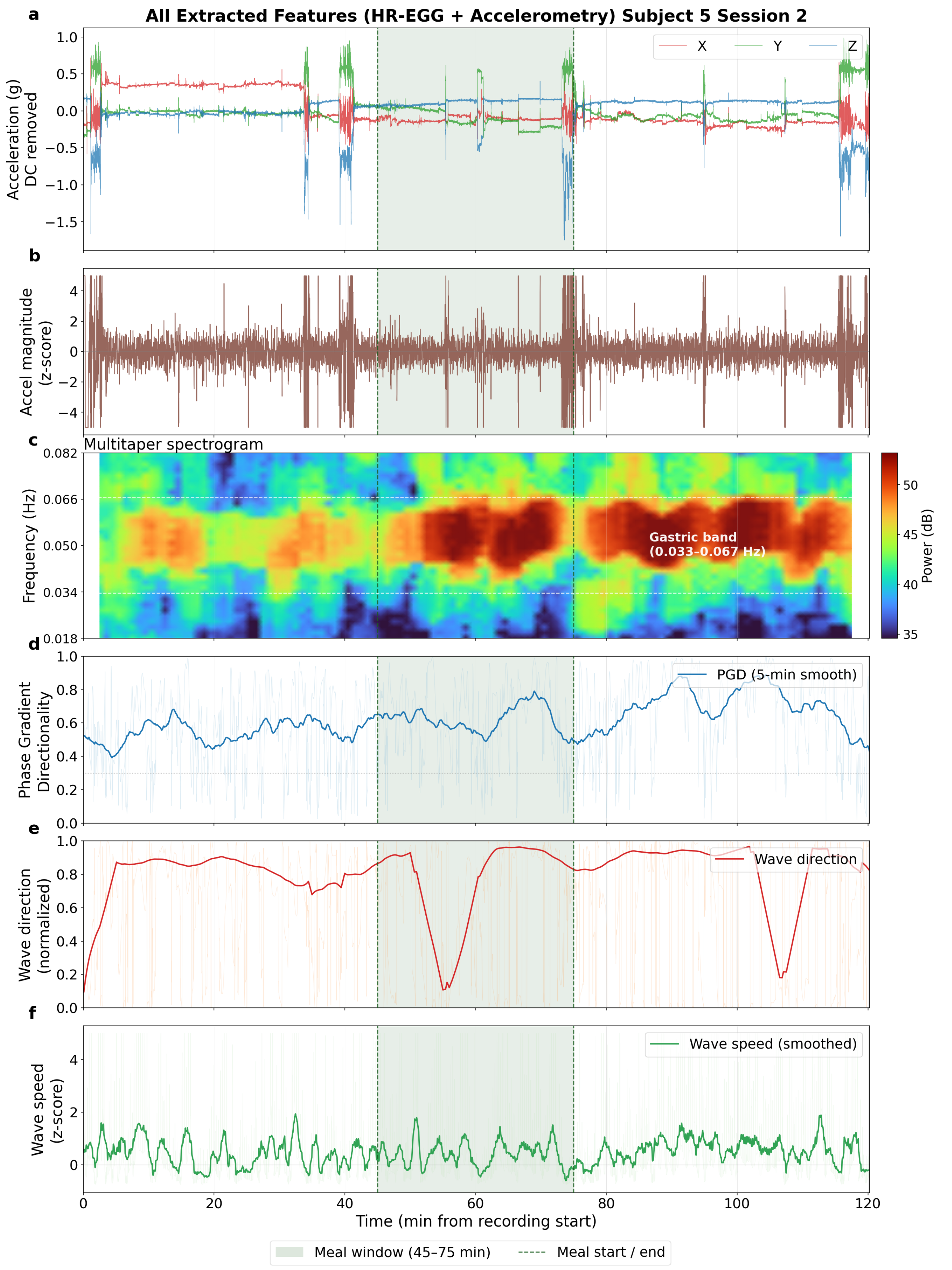


Figure S7. (Subject S5). (a) Triaxial accelerometry. (b) Accel magnitude z-score. (c) Multitaper HR-EGG spectrogram. (d) Phase gradient directionality (PGD). (e) Wave direction. (f) Wave speed.

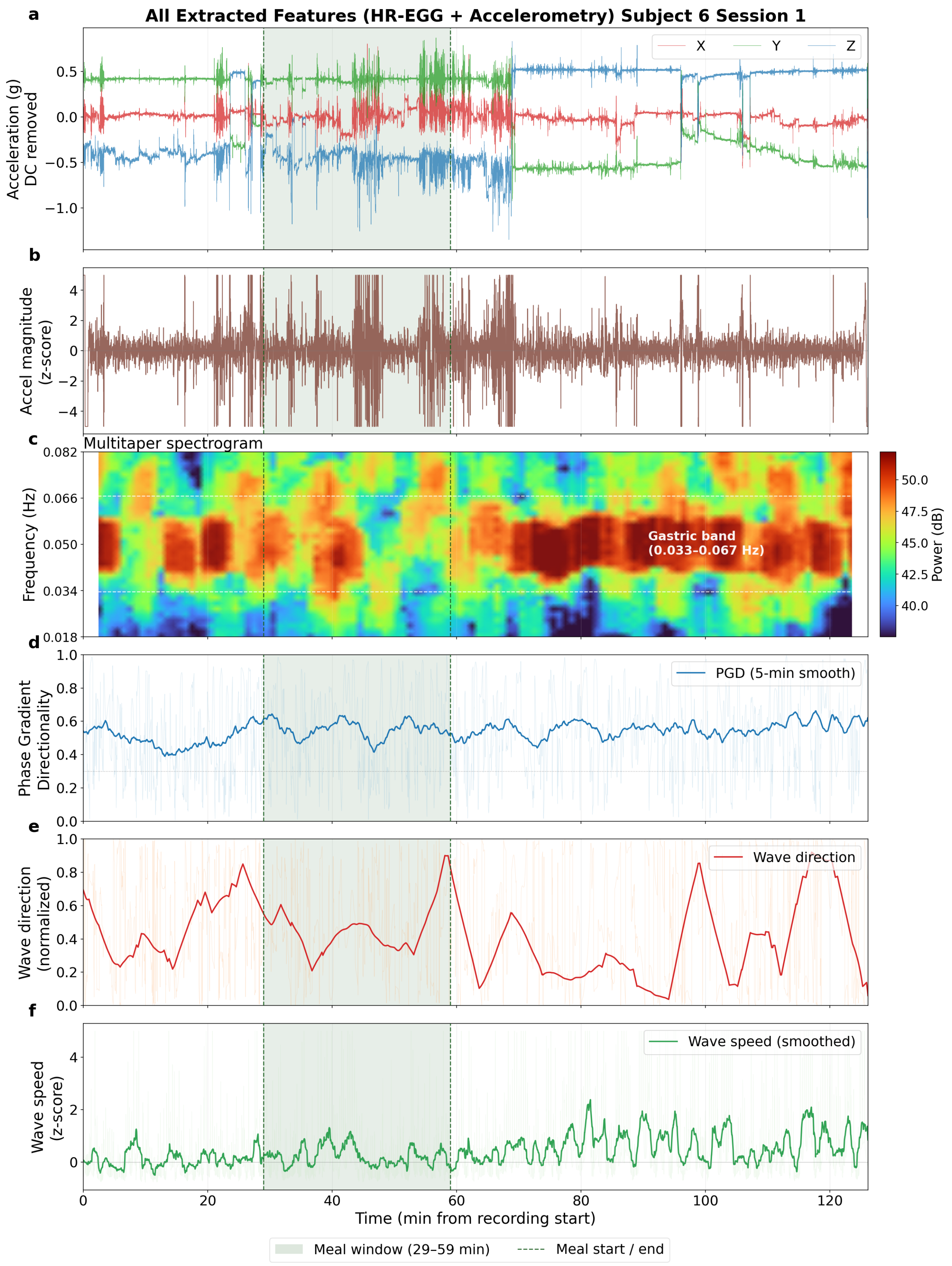


Figure S8. (Subject S6). (a) Triaxial accelerometry. (b) Accel magnitude z-score. (c) Multitaper HR-EGG spectrogram. (d) Phase gradient directionality (PGD). (e) Wave direction. (f) Wave speed.

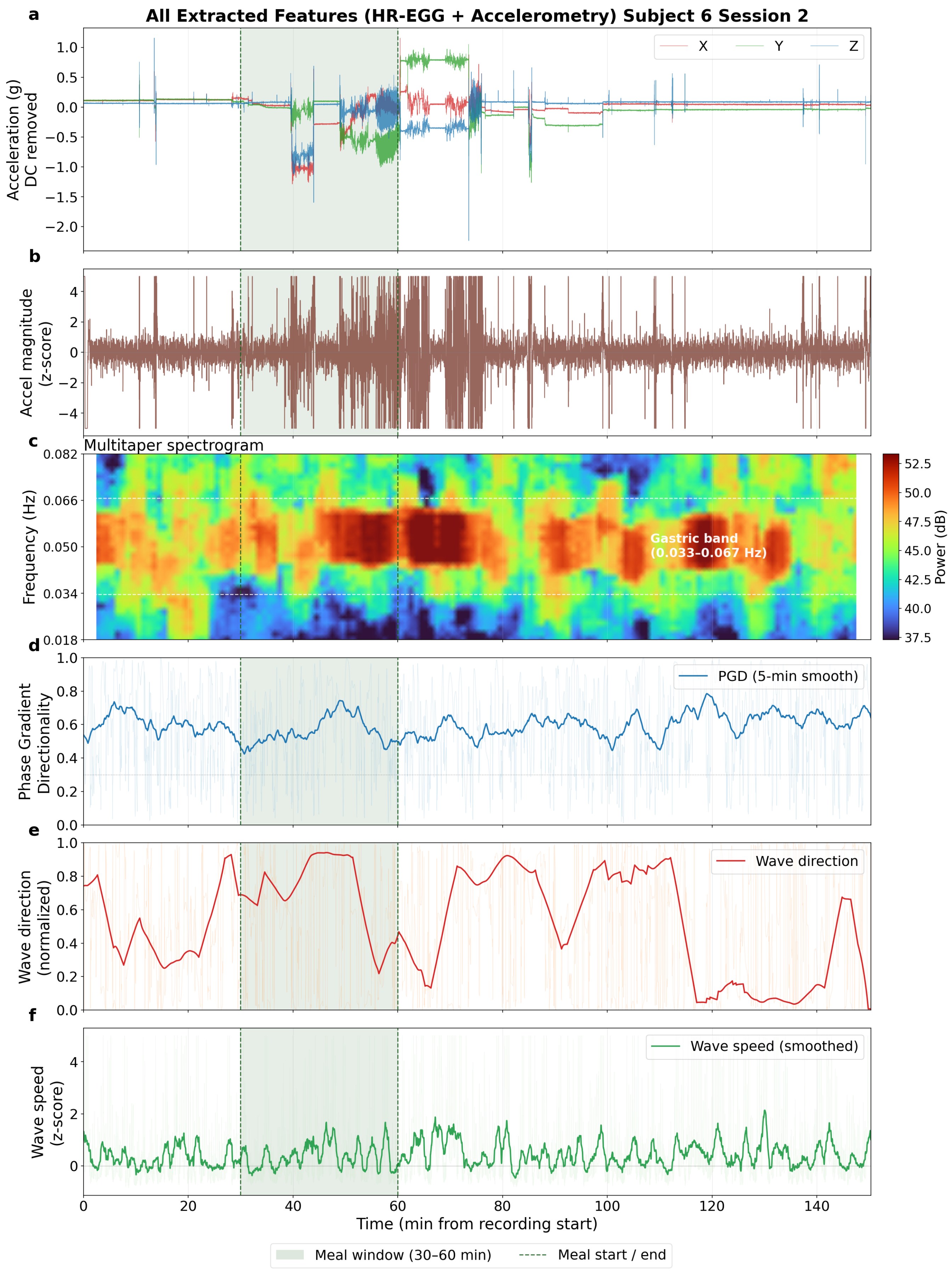


Figure S9. (Subject S6). (a) Triaxial accelerometry. (b) Accel magnitude z-score. (c) Multitaper HR-EGG spectrogram. (d) Phase gradient directionality (PGD). (e) Wave direction. (f) Wave speed.

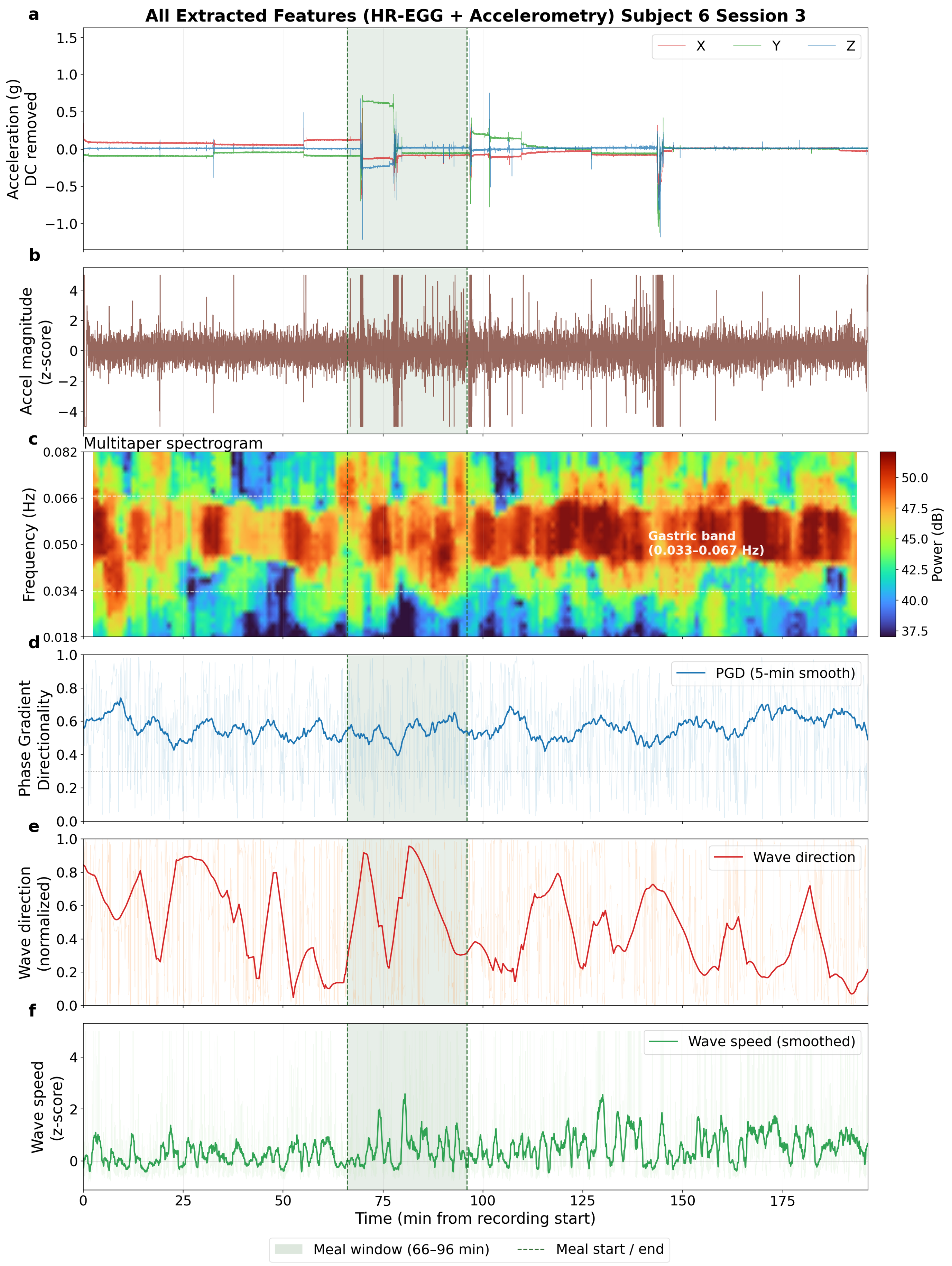


Figure S10. (Subject S6). (a) Triaxial accelerometry. (b) Accel magnitude z-score. (c) Multitaper HR-EGG spectrogram. (d) Phase gradient directionality (PGD). (e) Wave direction. (f) Wave speed.

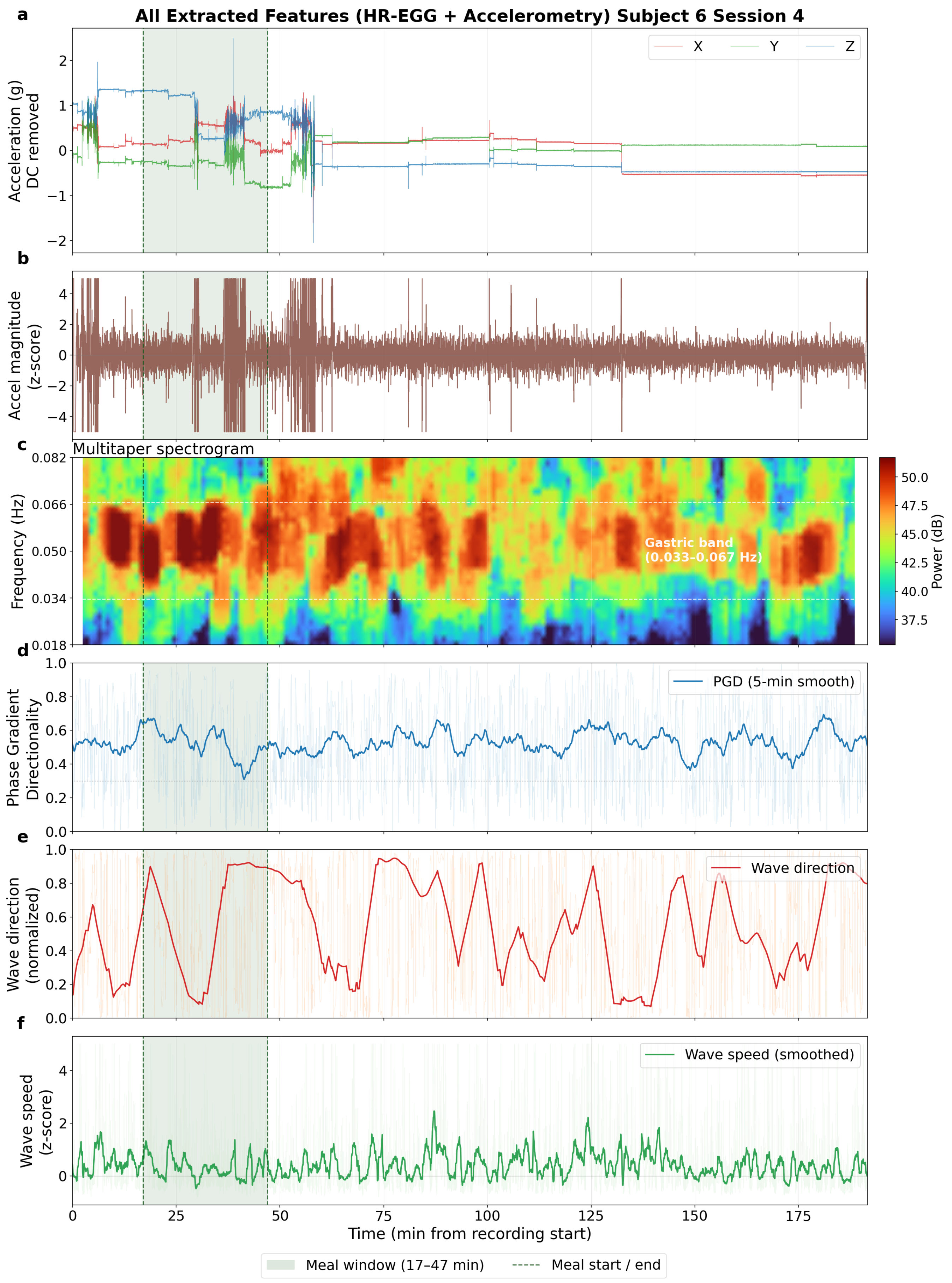


Figure S11. (Subject S6). (a) Triaxial accelerometry. (b) Accel magnitude z-score. (c) Multitaper HR-EGG spectrogram. (d) Phase gradient directionality (PGD). (e) Wave direction. (f) Wave speed.

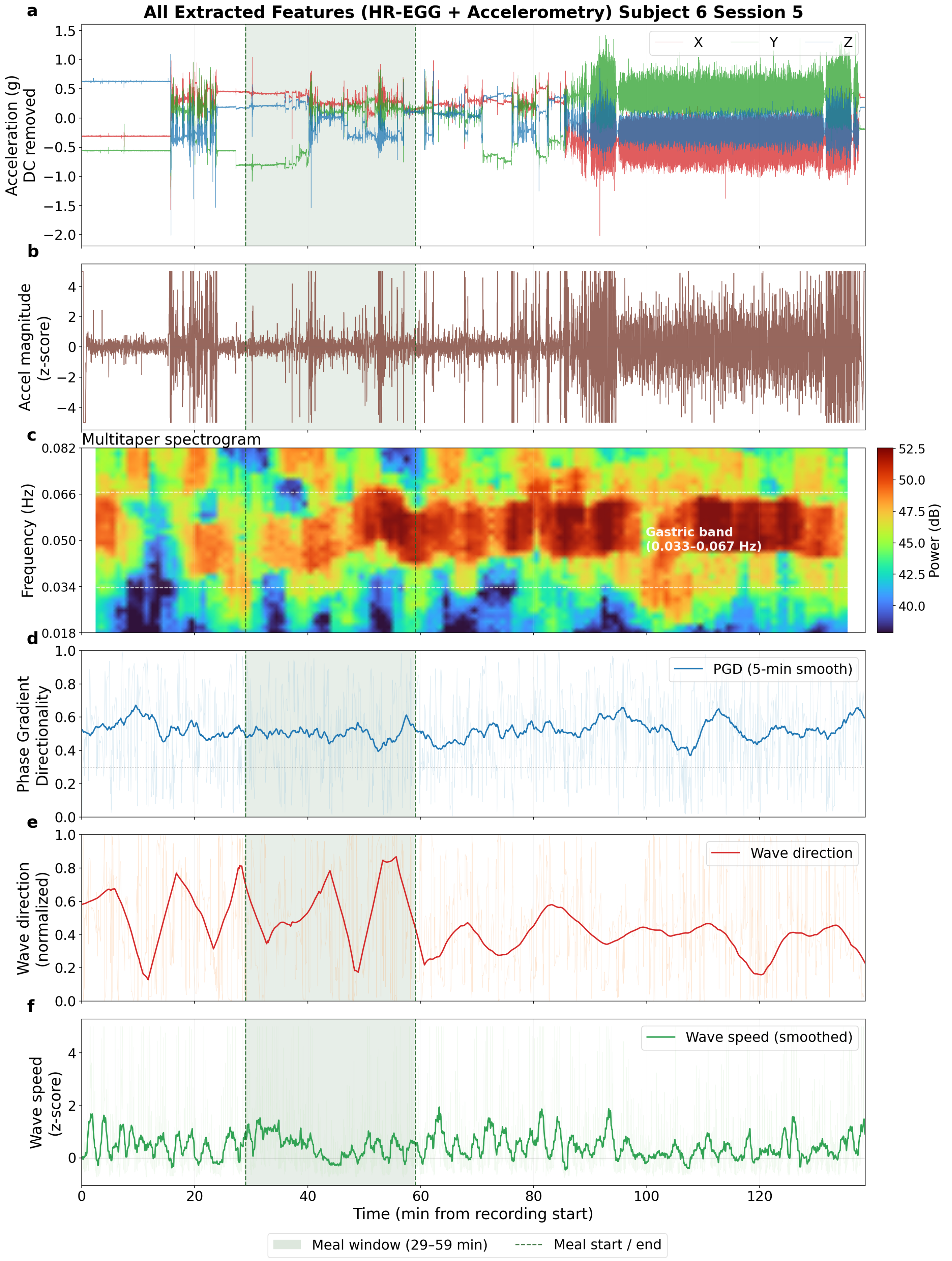


Figure S12. (Subject S6). (a) Triaxial accelerometry. (b) Accel magnitude z-score. (c) Multitaper HR-EGG spectrogram. (d) Phase gradient directionality (PGD). (e) Wave direction. (f) Wave speed.

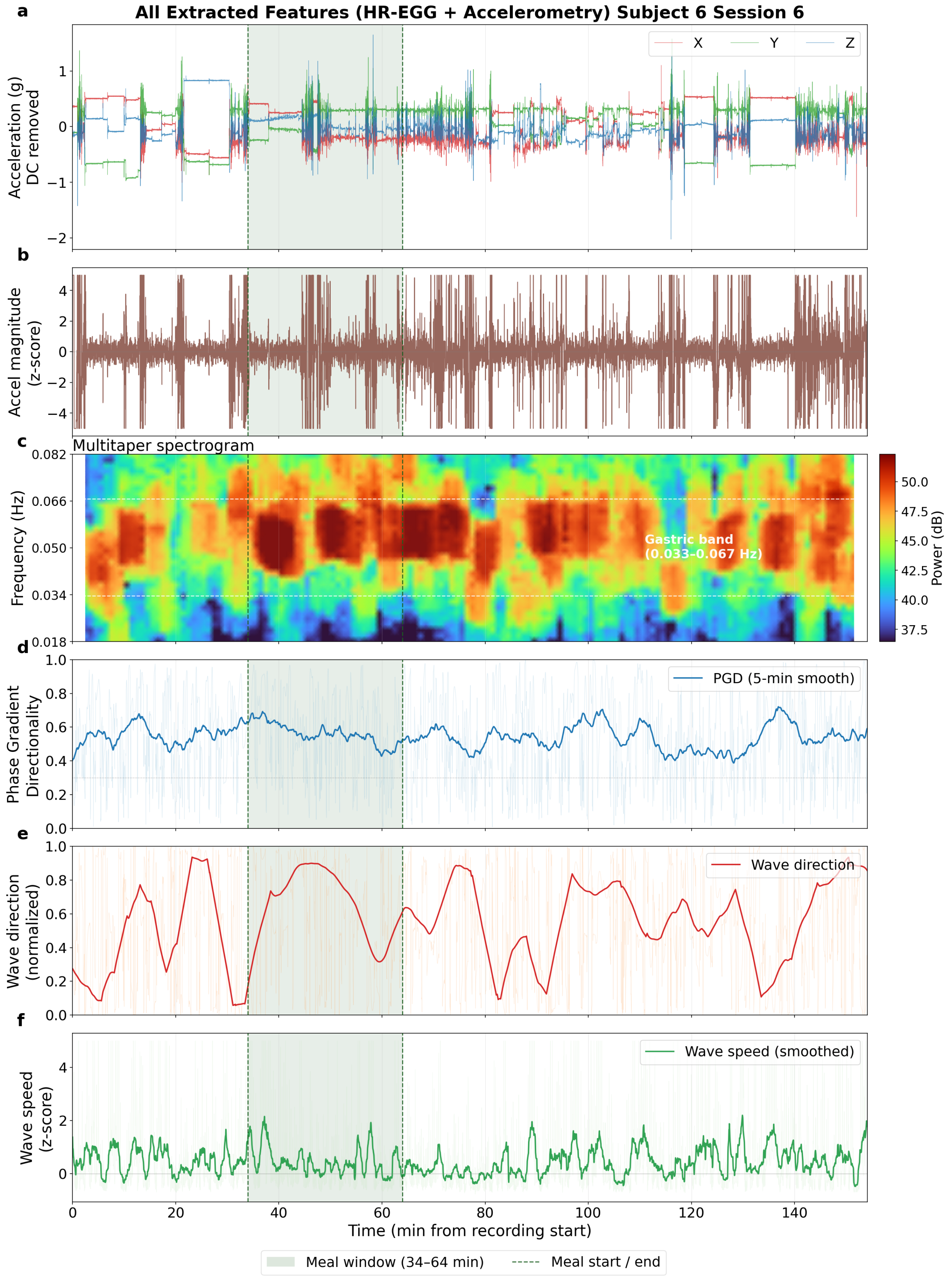


Figure S13. (Subject S6). (a) Triaxial accelerometry. (b) Accel magnitude z-score. (c) Multitaper HR-EGG spectrogram. (d) Phase gradient directionality (PGD). (e) Wave direction. (f) Wave speed.

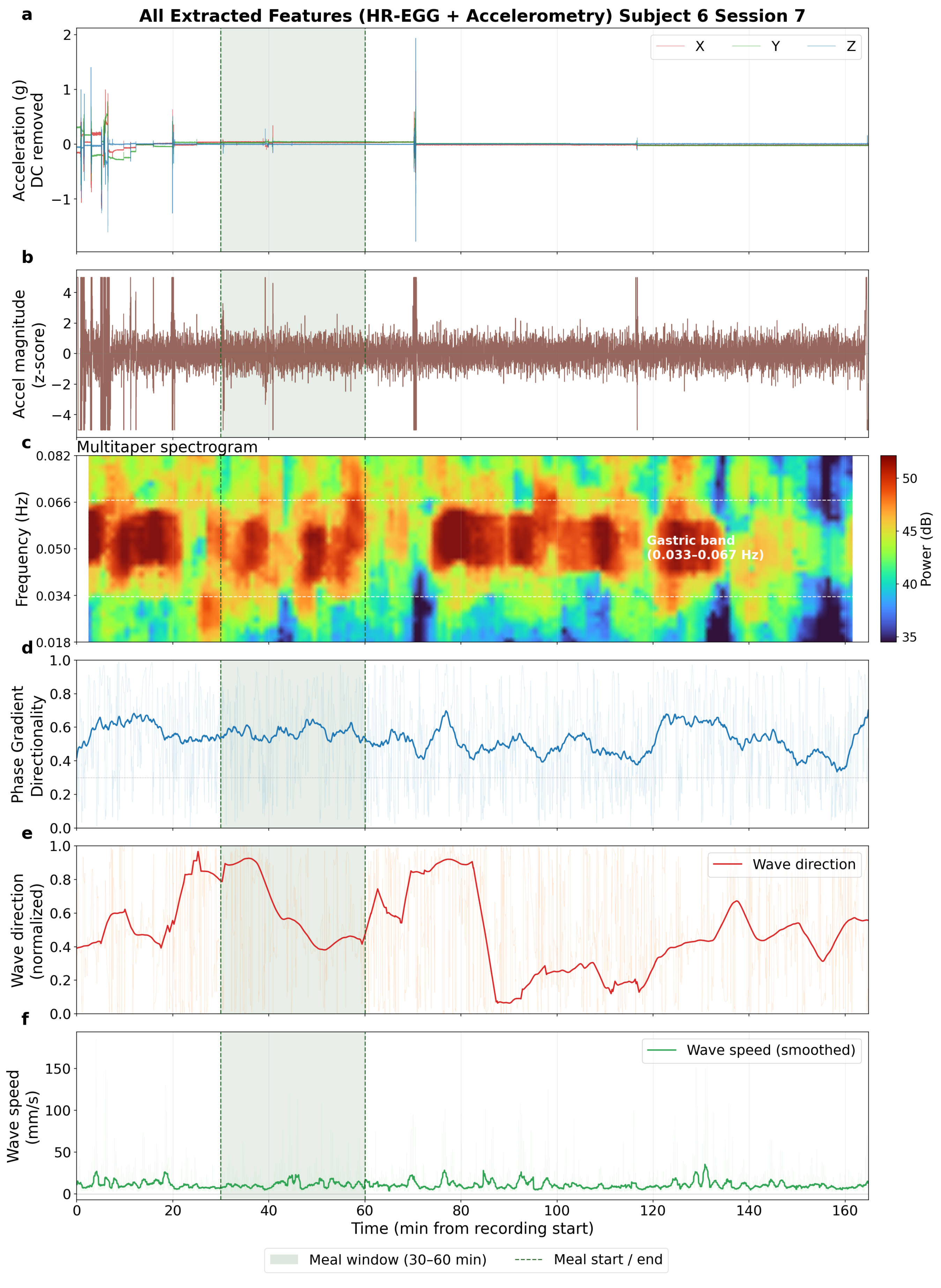


Figure S14. (Subject S6). (a) Triaxial accelerometry. (b) Accel magnitude z-score. (c) Multitaper HR-EGG spectrogram. (d) Phase gradient directionality (PGD). (e) Wave direction. (f) Wave speed.

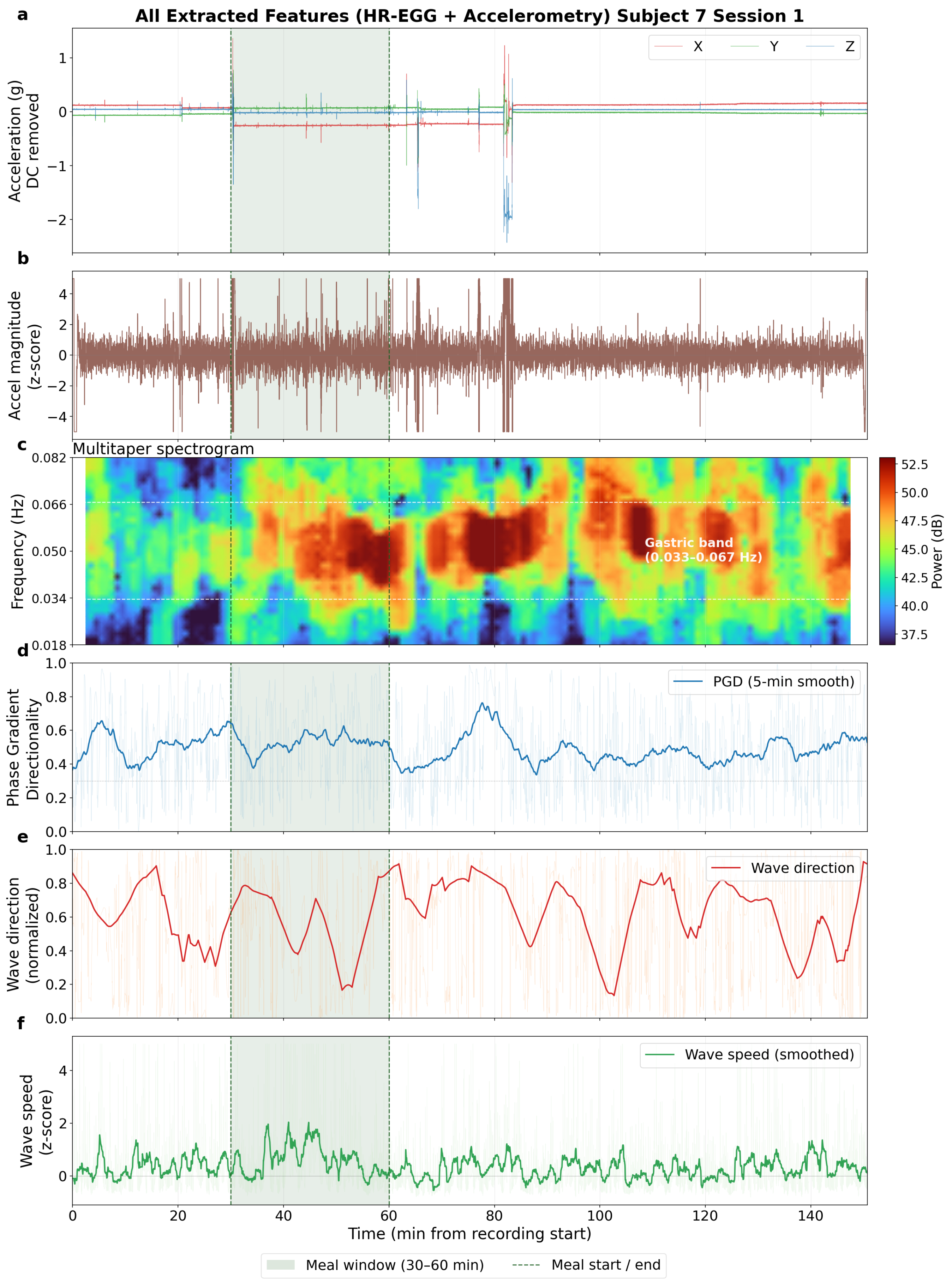


Figure S15. (Subject S7). (a) Triaxial accelerometry. (b) Accel magnitude z-score. (c) Multitaper HR-EGG spectrogram. (d) Phase gradient directionality (PGD). (e) Wave direction. (f) Wave speed.

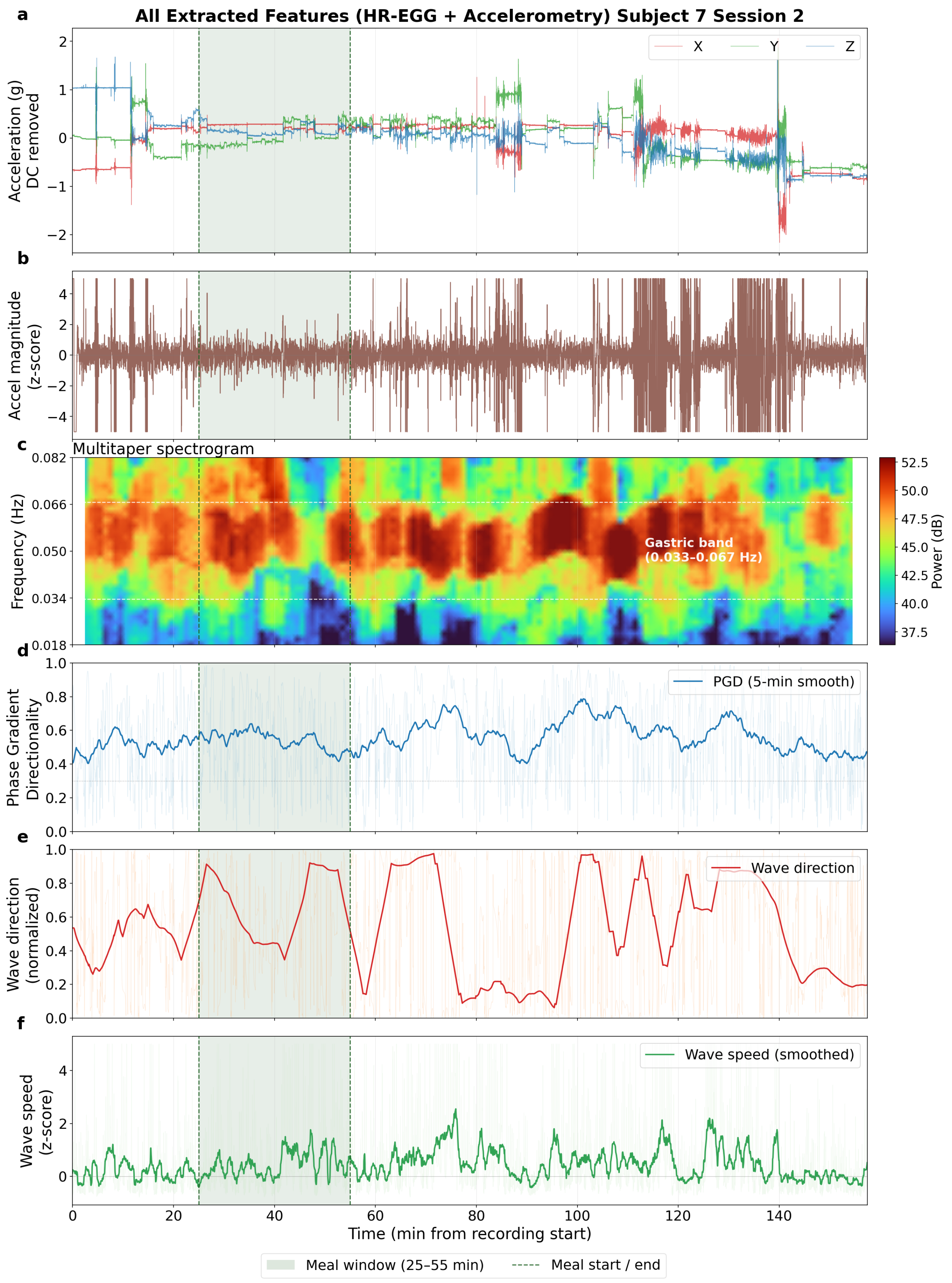


Figure S16. (Subject S7). (a) Triaxial accelerometry. (b) Accel magnitude z-score. (c) Multitaper HR-EGG spectrogram. (d) Phase gradient directionality (PGD). (e) Wave direction. (f) Wave speed.

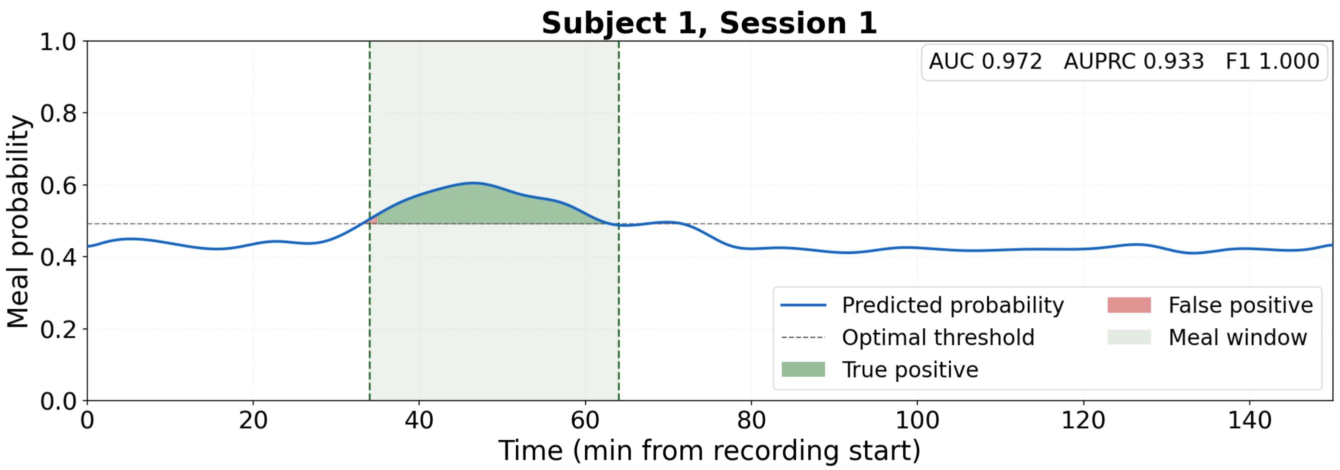


Figure S17. S1, Session 1. LOSO-subject prediction probability over time. AUROC=0.972, AUPRC=0.933, F1=1.000.

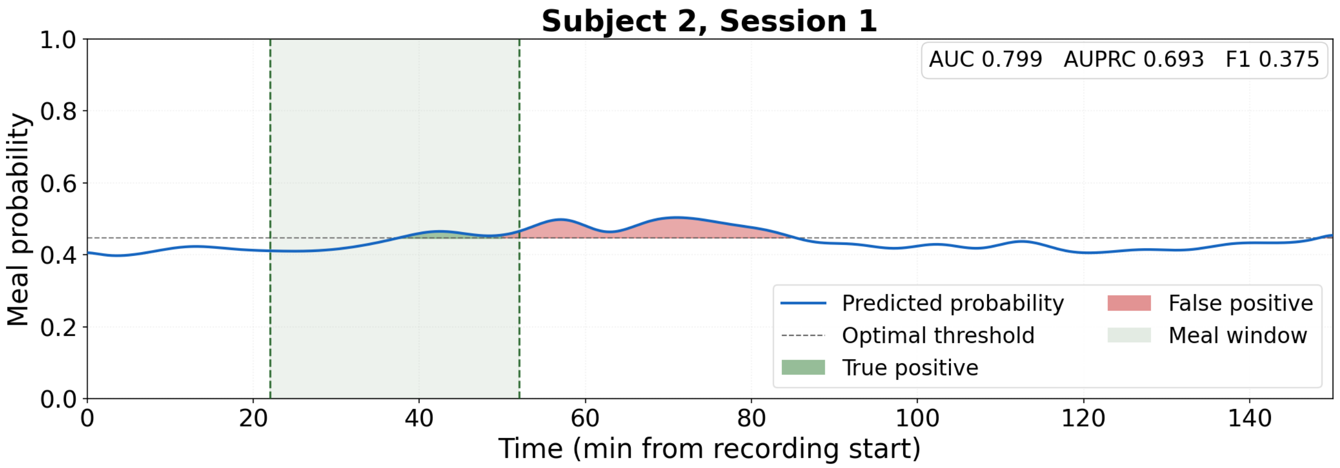


Figure S18. S2, Session 1. LOSO-subject prediction probability over time. AUROC=0.799, AUPRC=0.693, F1=0.375.

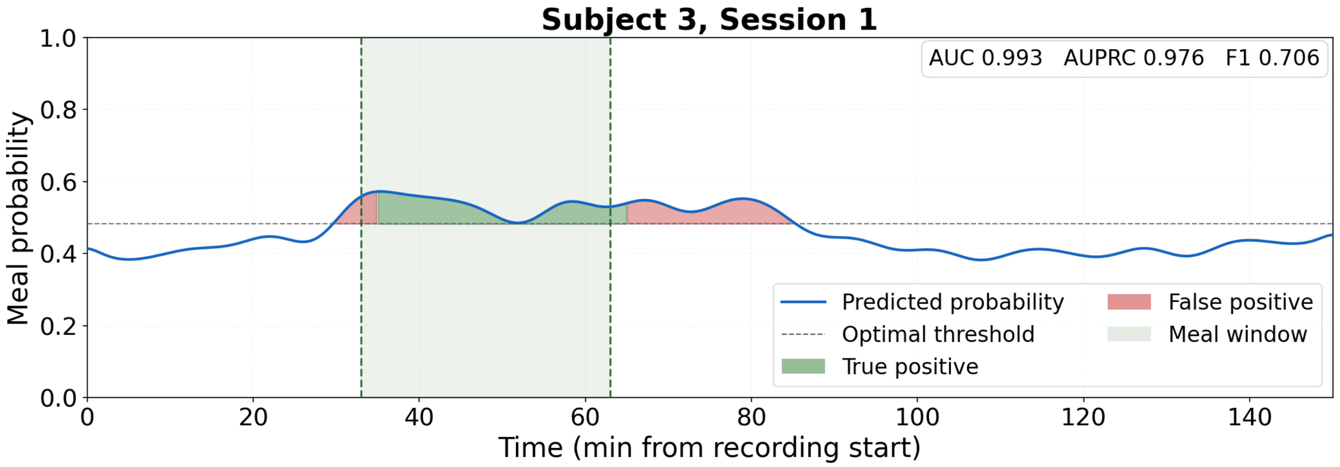


Figure S19. S3, Session 1. LOSO-subject prediction probability over time. AUROC=0.993, AUPRC=0.976, F1=0.706.

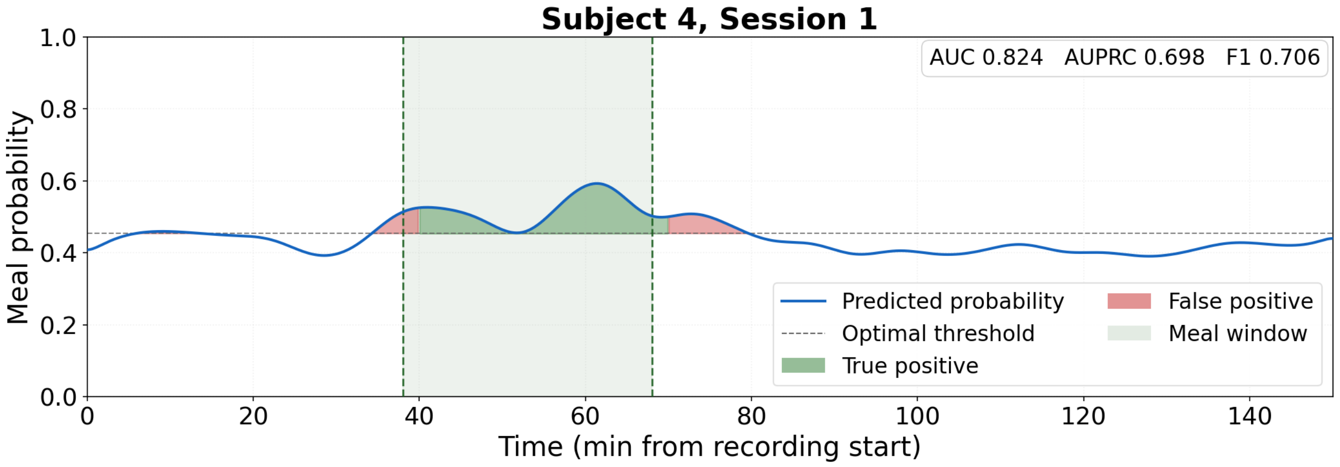


Figure S20. S4, Session 1. LOSO-subject prediction probability over time. AUROC=0.824, AUPRC=0.698, F1=0.706.

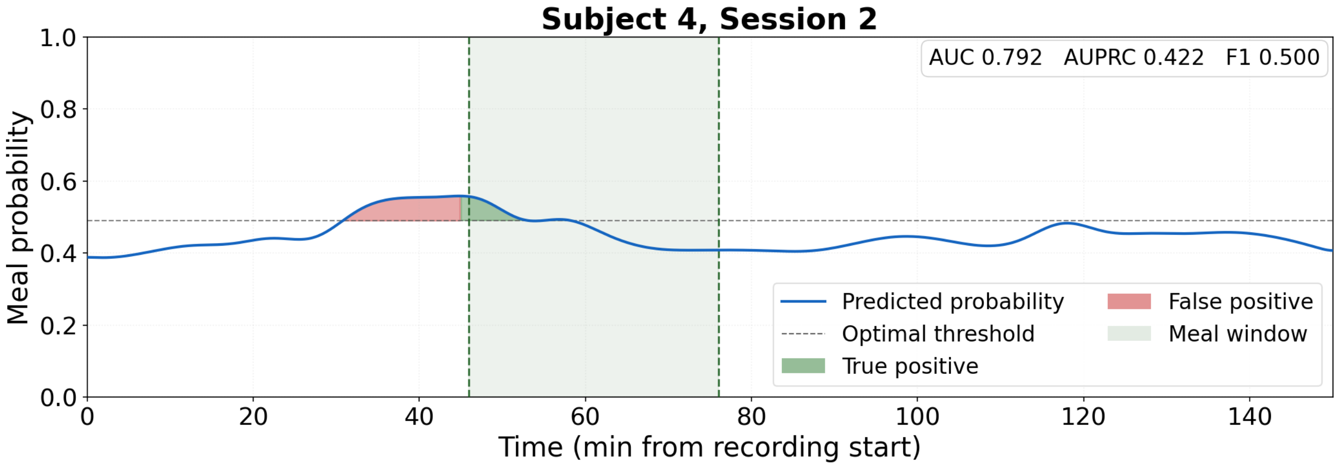


Figure S21. S4, Session 2. LOSO-subject prediction probability over time. AUROC=0.792, AUPRC=0.422, F1=0.500.

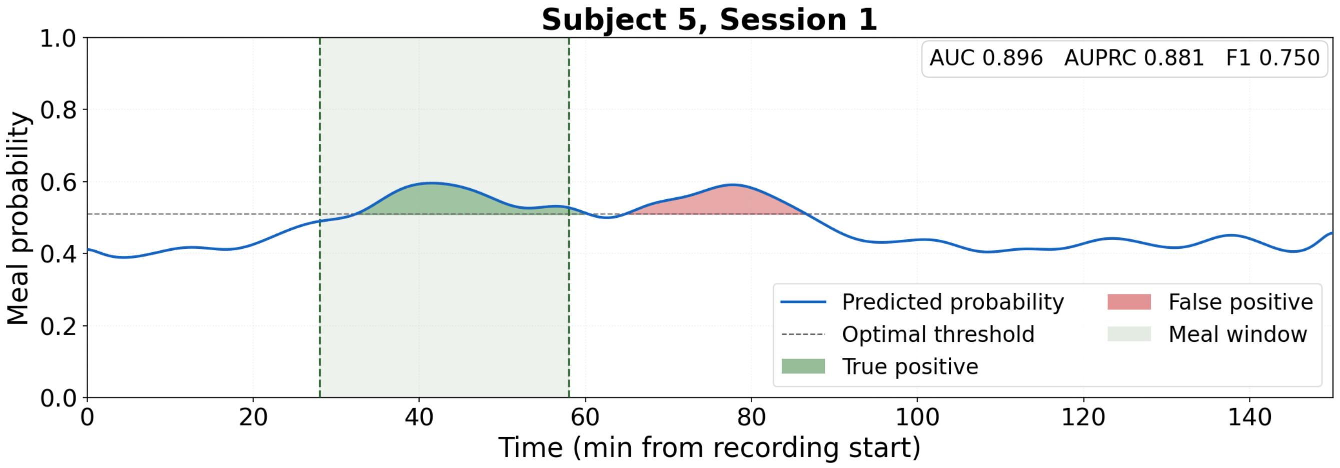


Figure S22. S5, Session 1. LOSO-subject prediction probability over time. AUROC=0.896, AUPRC=0.881, F1=0.750.

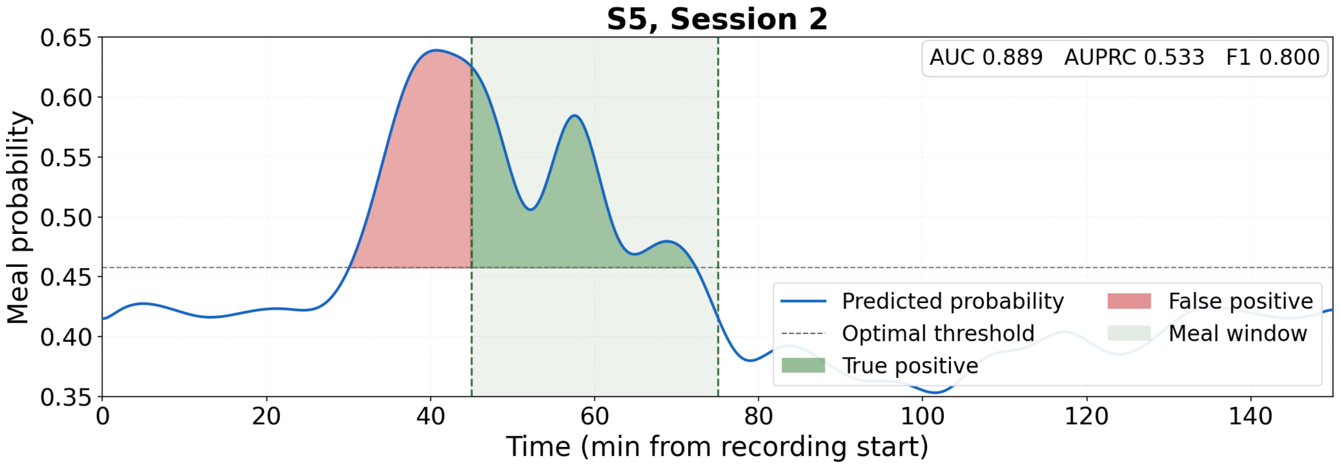


Figure S23. S5, Session 2. LOSO-subject prediction probability over time. AUROC=0.889, AUPRC=0.533, F1=0.800.

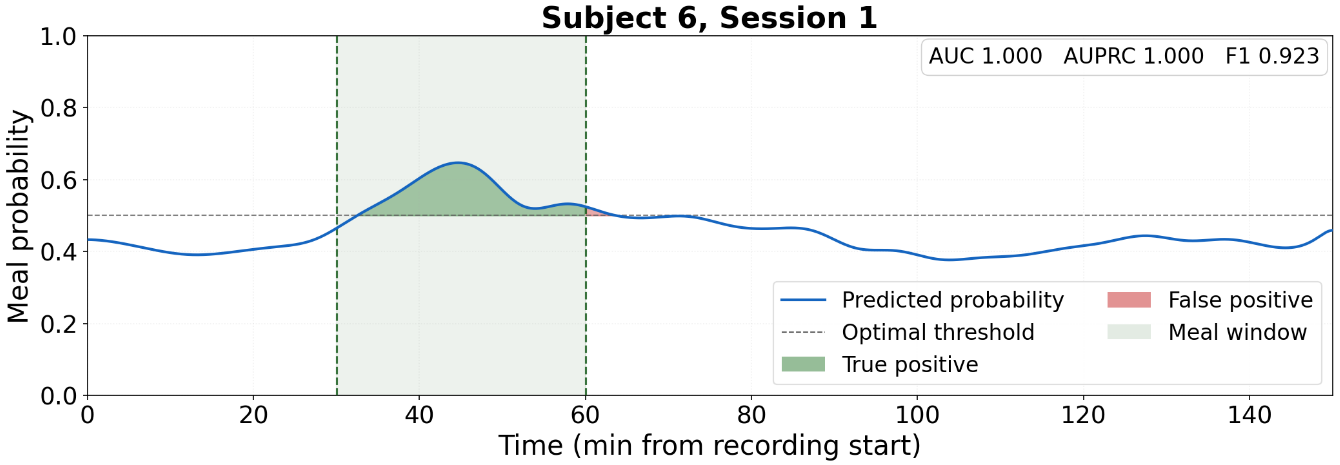


Figure S24. S6, Session 1. LOSO-subject prediction probability over time. AUROC=1.000, AUPRC=1.000, F1=0.923.

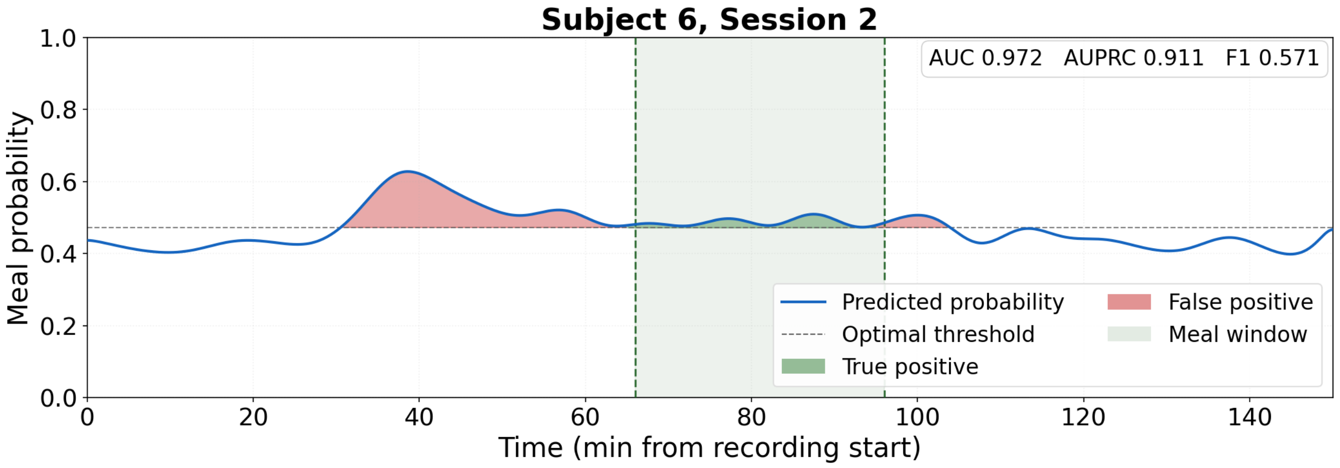


Figure S25. S6, Session 2. LOSO-subject prediction probability over time. AUROC=0.972, AUPRC=0.911, F1=0.571.

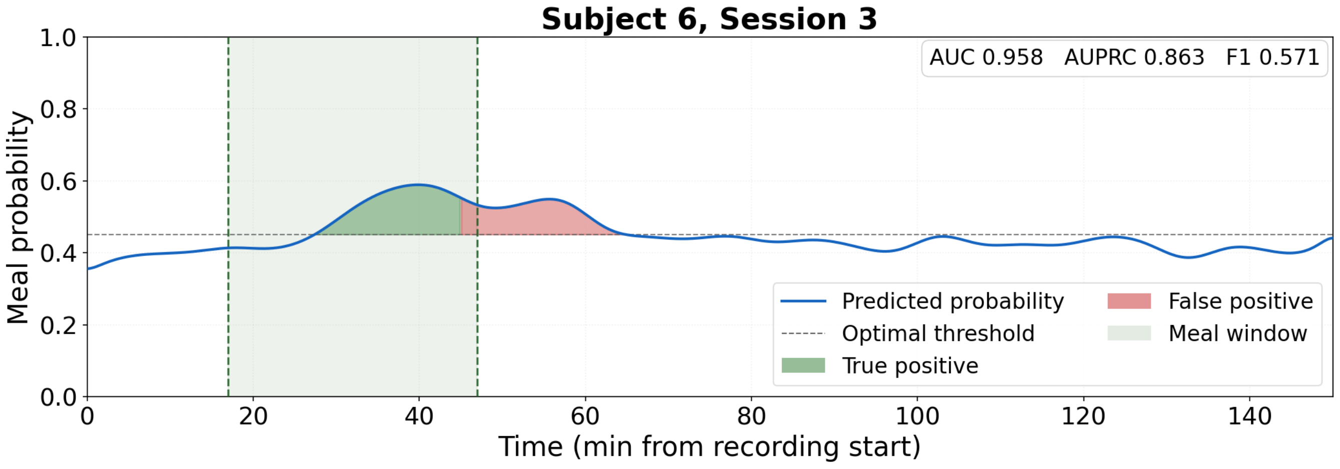


Figure S26. S6, Session 3. (LOSO-subject prediction probability over time. AUROC=0.944, AUPRC=0.862, F1=0.706.

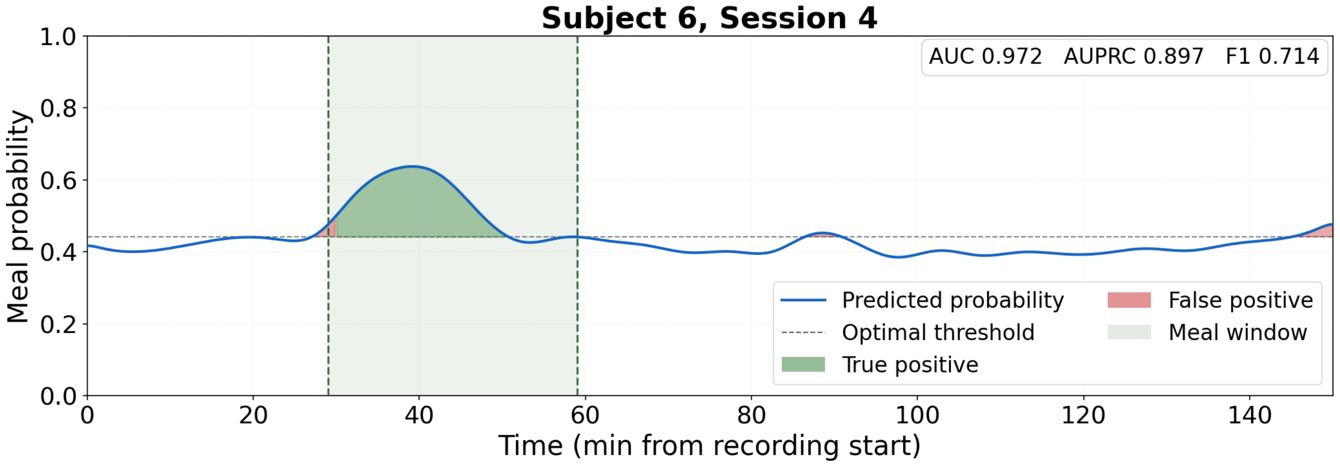


Figure S27. S6, Session 4. LOSO-subject prediction probability over time. AUROC=0.972, AUPRC=0.897, F1=0.714.

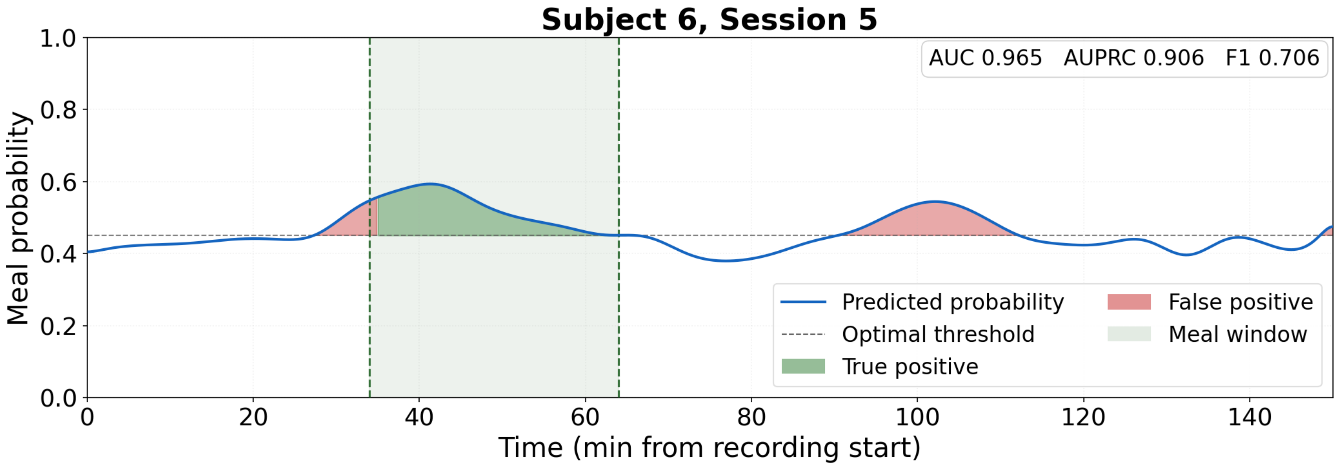


Figure S28. S6, Session 5. LOSO-subject prediction probability over time. AUROC=0.965, AUPRC=0.906, F1=0.706

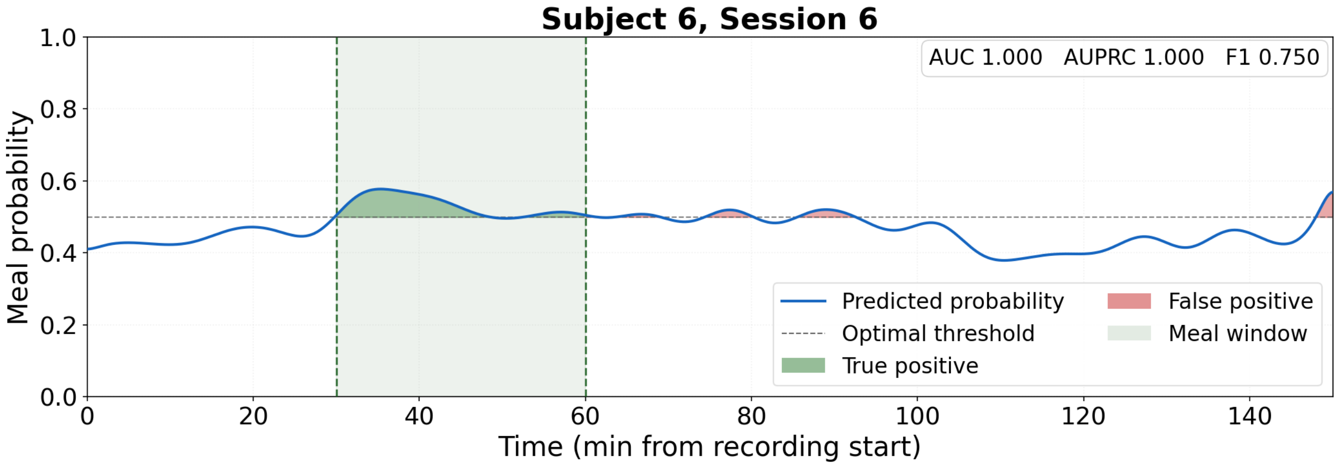


Figure S29. S6, Session 6. LOSO-subject prediction probability over time. AUROC=1.000, AUPRC=1.000, F1=0.750.

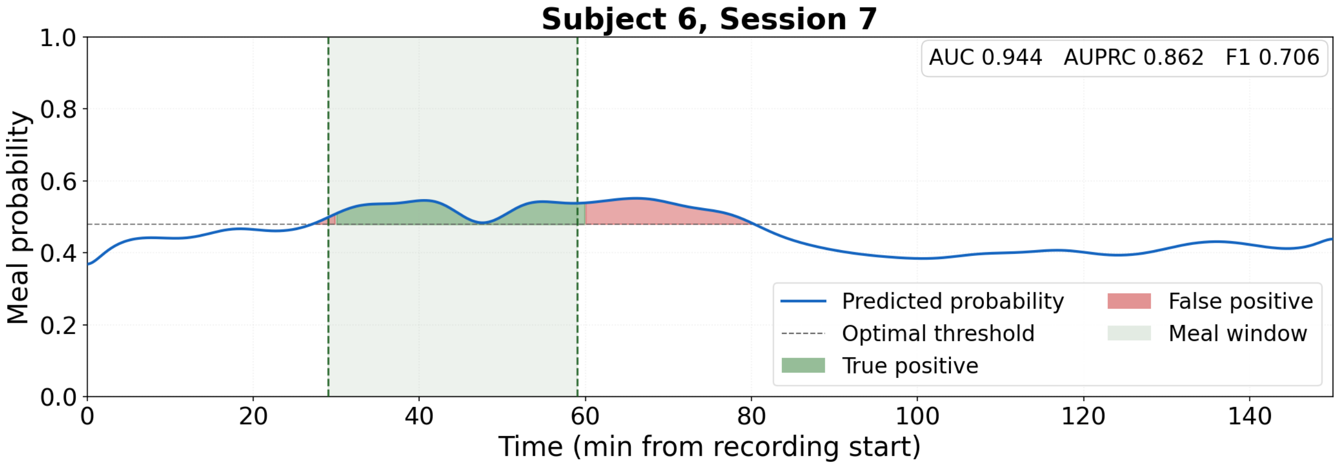


Figure S30. S6, Session 7. LOSO-subject prediction probability over time. AUROC=0.944, AUPRC=0.862, F1=0.706.

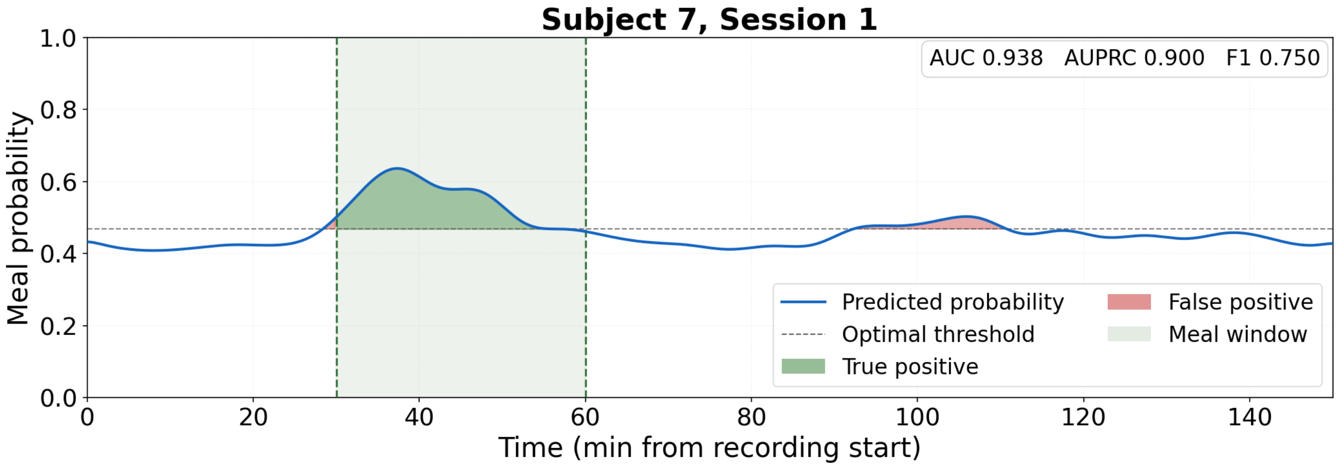


Figure S31. S7, Session 1. LOSO-subject prediction probability over time. AUROC=0.938, AUPRC=0.900, F1=0.750.

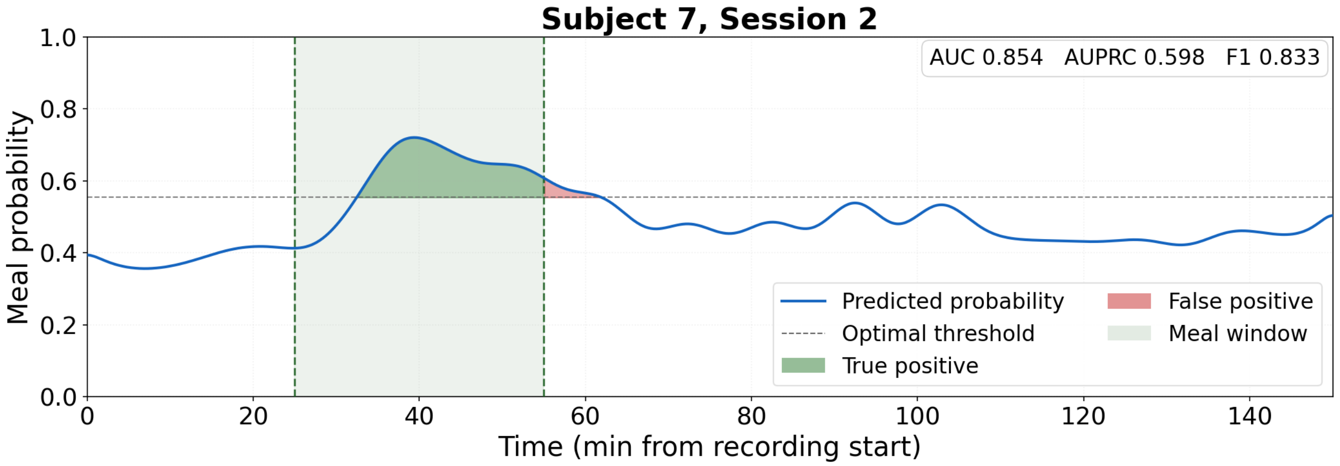


Figure S32. S7, Session 2. LOSO-subject prediction probability over time. AUROC=0.854, AUPRC=0.598, F1=0.833.

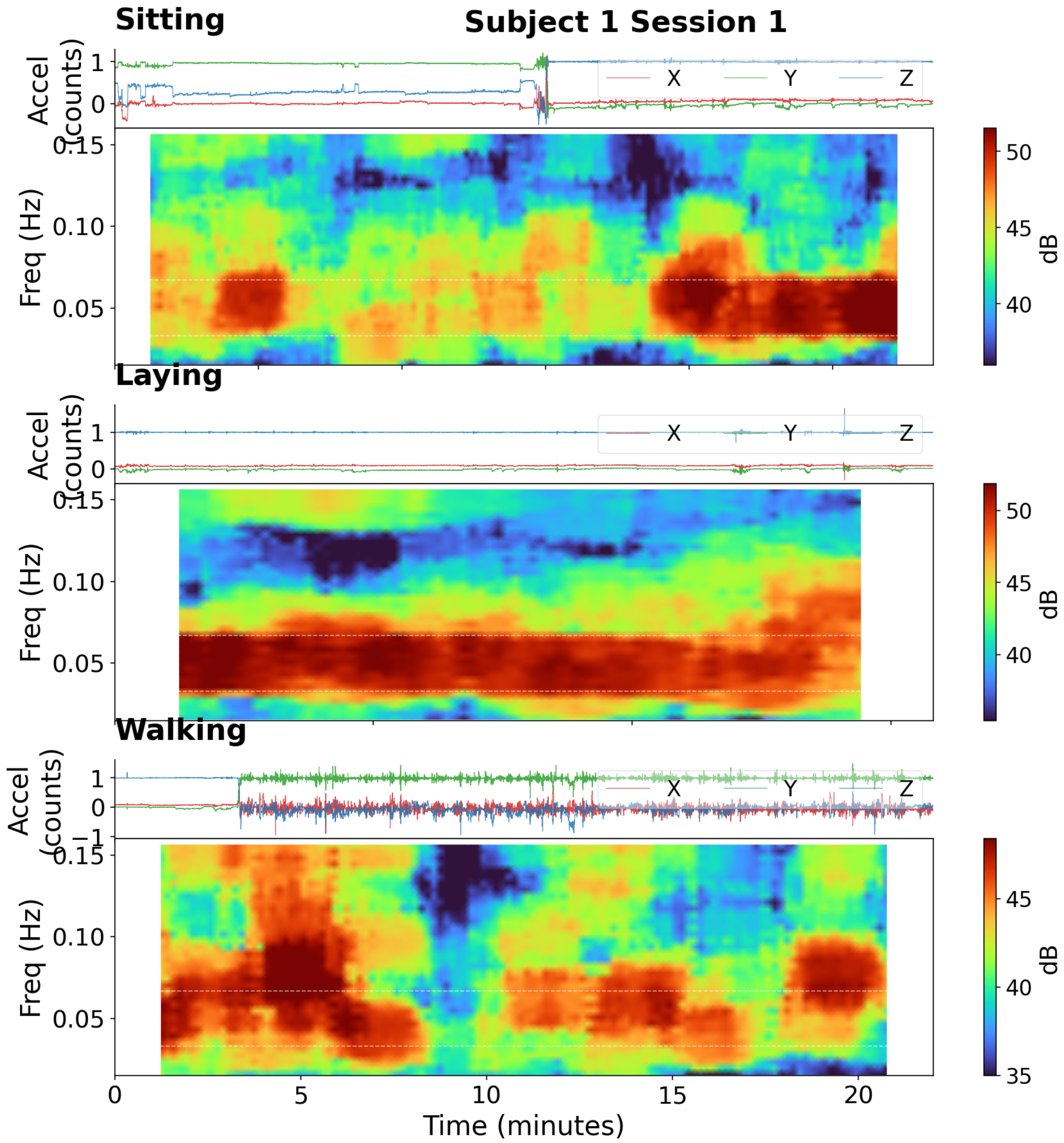


Figure S33. (Subject S1). Post-meal activity phases: sitting, laying, walking. Top panel: triaxial accelerometry. Bottom panel: multitaper HR-EGG spectrogram (120-s window, 15-s step). Dashed white lines mark the 2–4 cpm normogastric band.

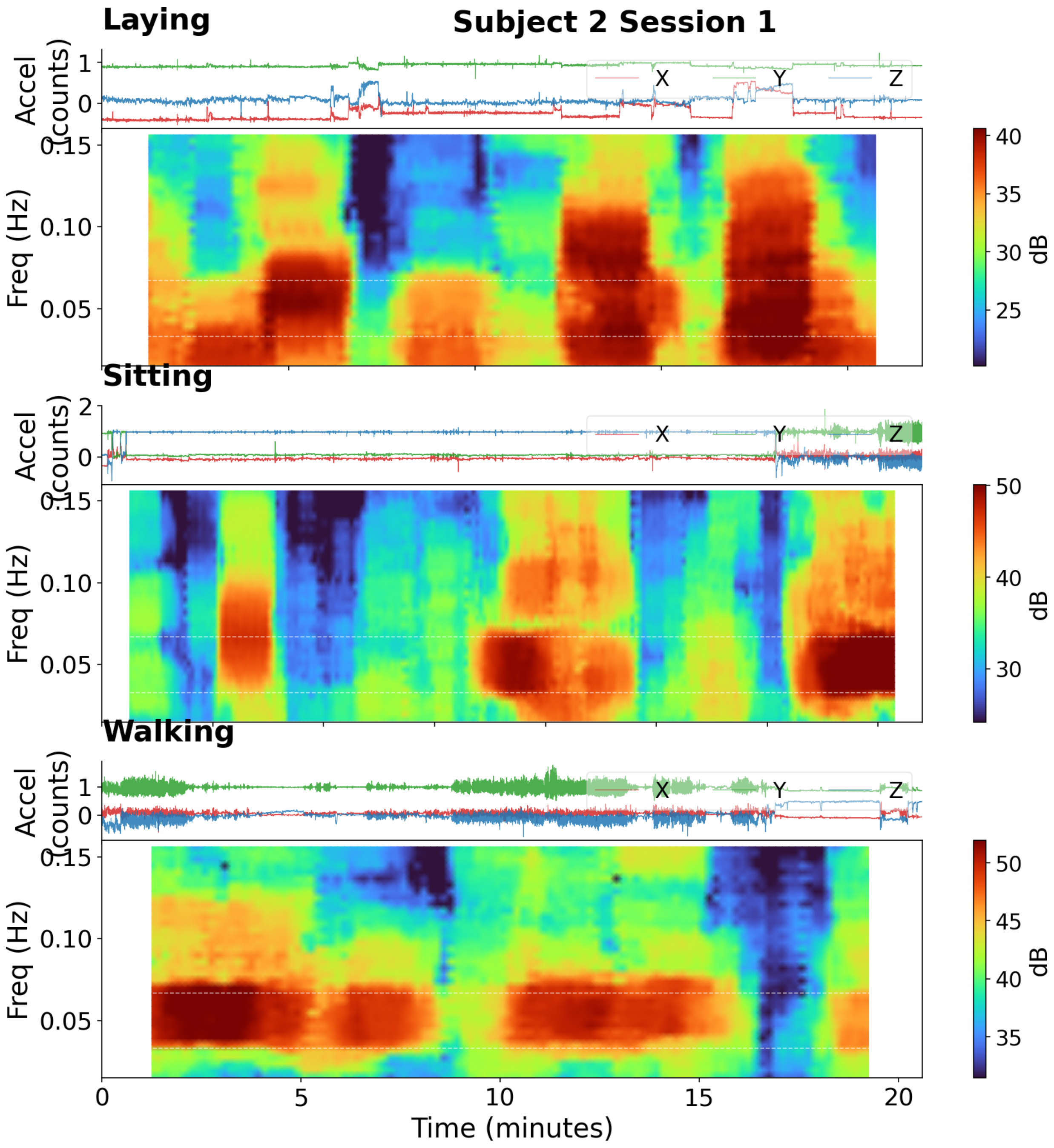


Figure S34. (Subject S2). Post-meal activity phases: laying, sitting, walking. Top panel: triaxial accelerometry. Bottom panel: multitaper HR-EGG spectrogram (120-s window, 15-s step). Dashed white lines mark the 2–4 cpm normogastric band.

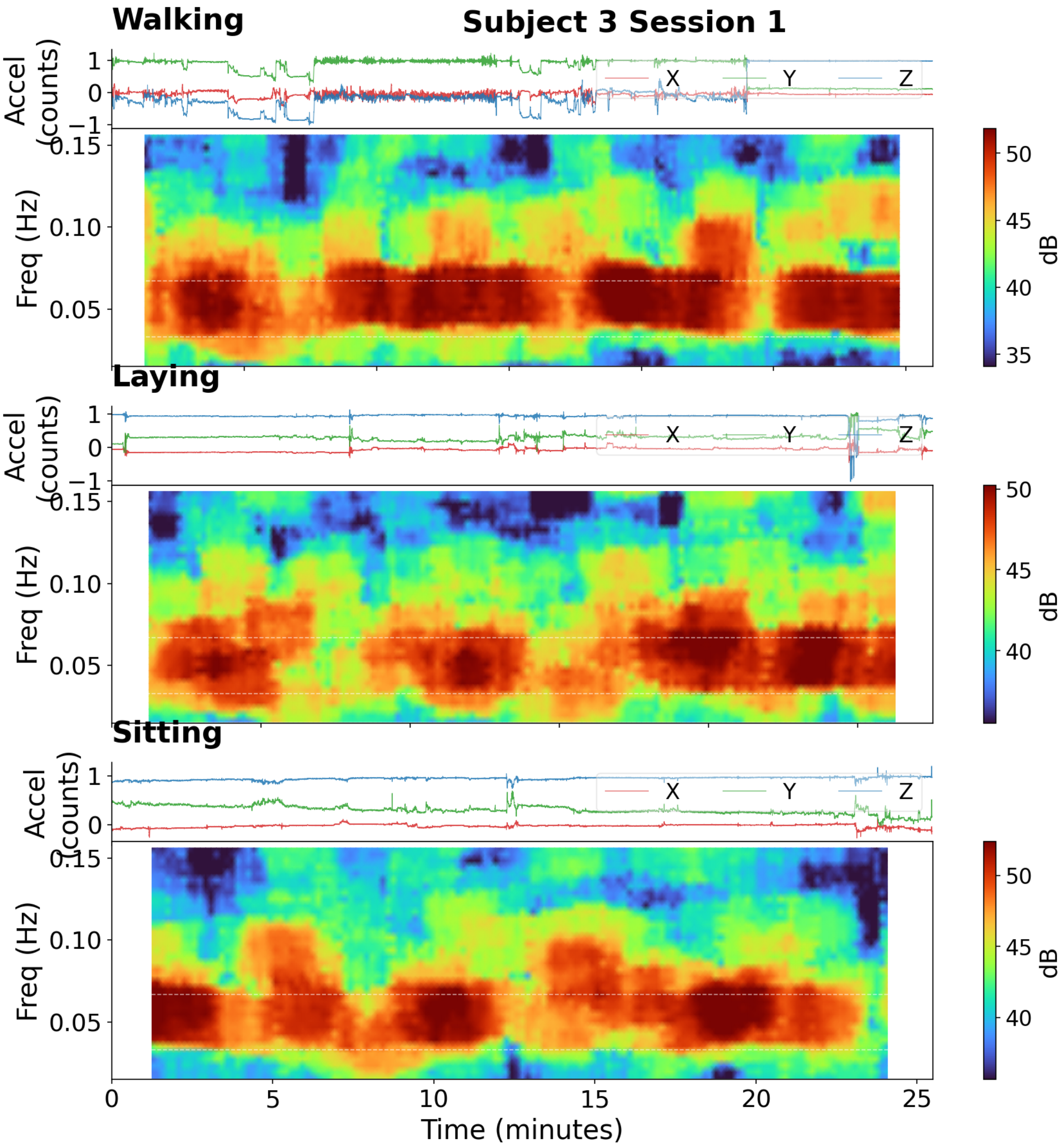


Figure S35. (Subject S3). Post-meal activity phases: walking, sitting, laying. Top panel: triaxial accelerometry. Bottom panel: multitaper HR-EGG spectrogram (120-s window, 15-s step). Dashed white lines mark the 2–4 cpm normogastric band.

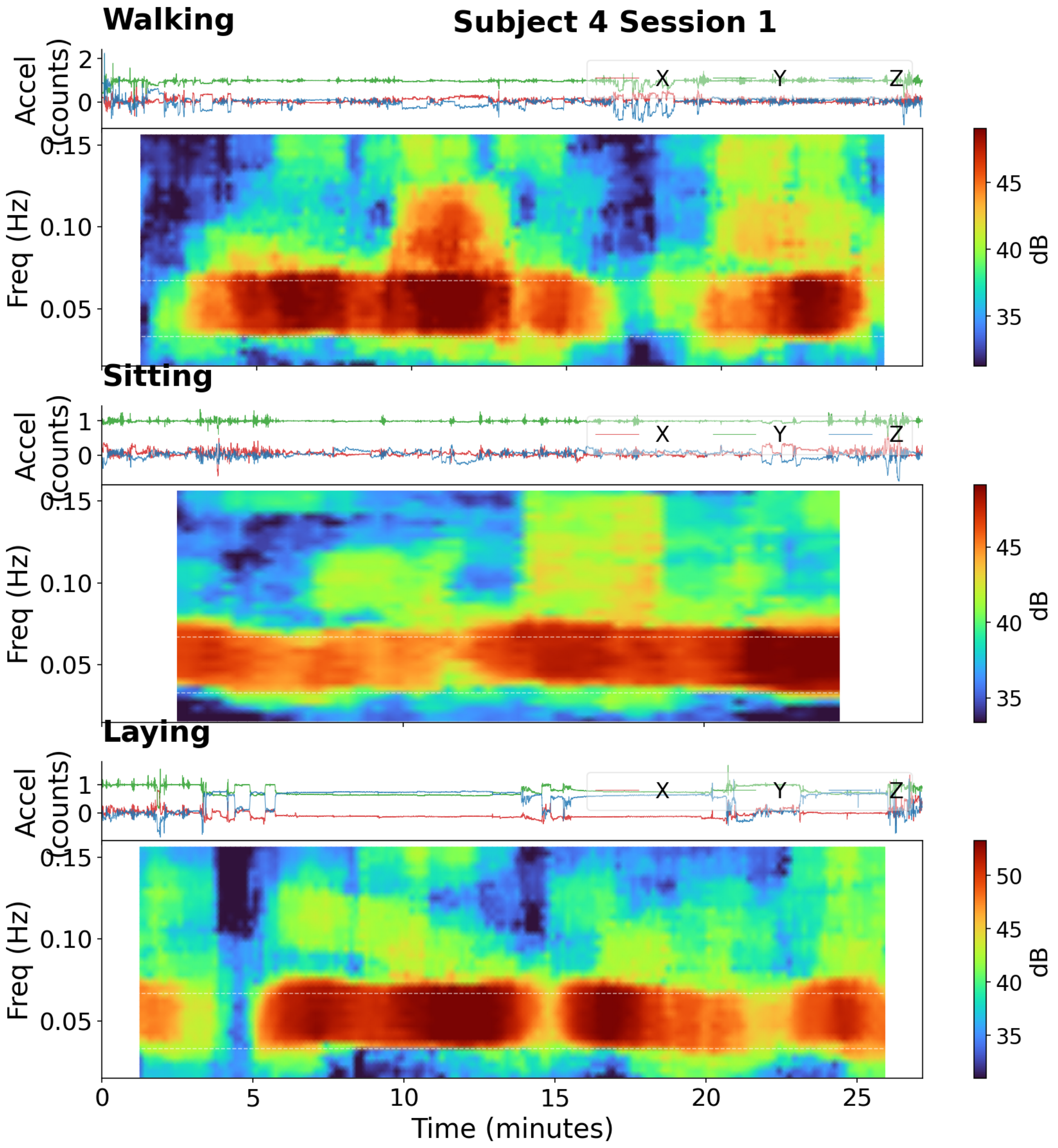


Figure S36. (Subject S4). Post-meal activity phases: walking, sitting, laying. Top panel: triaxial accelerometry. Bottom panel: multitaper HR-EGG spectrogram (120-s window, 15-s step). Dashed white lines mark the 2–4 cpm normogastric band.

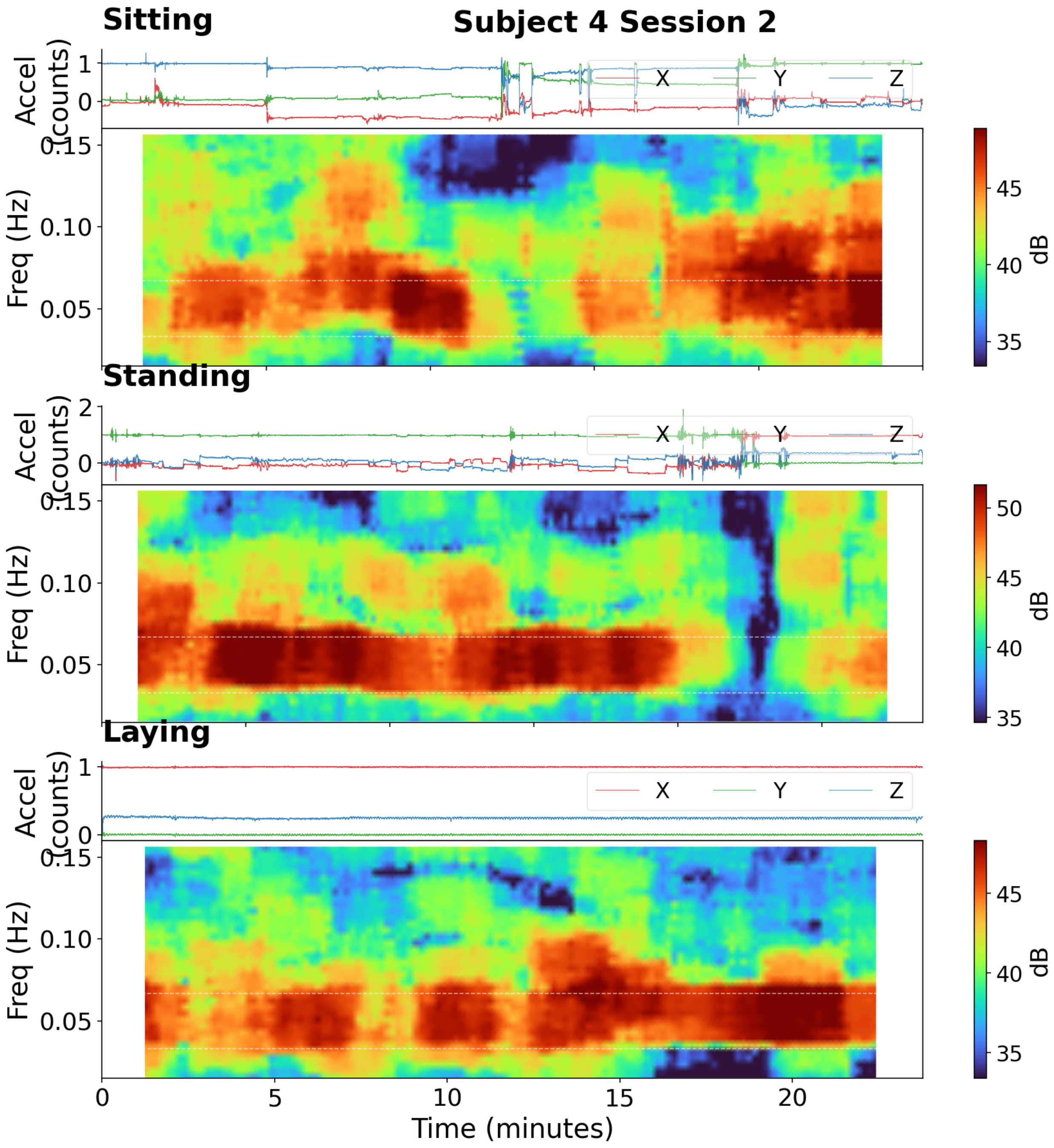


Figure S37. (Subject S4). Post-meal activity phases: sitting, standing, laying. Top panel: triaxial accelerometry. Bottom panel: multitaper HR-EGG spectrogram (120-s window, 15-s step). Dashed white lines mark the 2–4 cpm normogastric band.

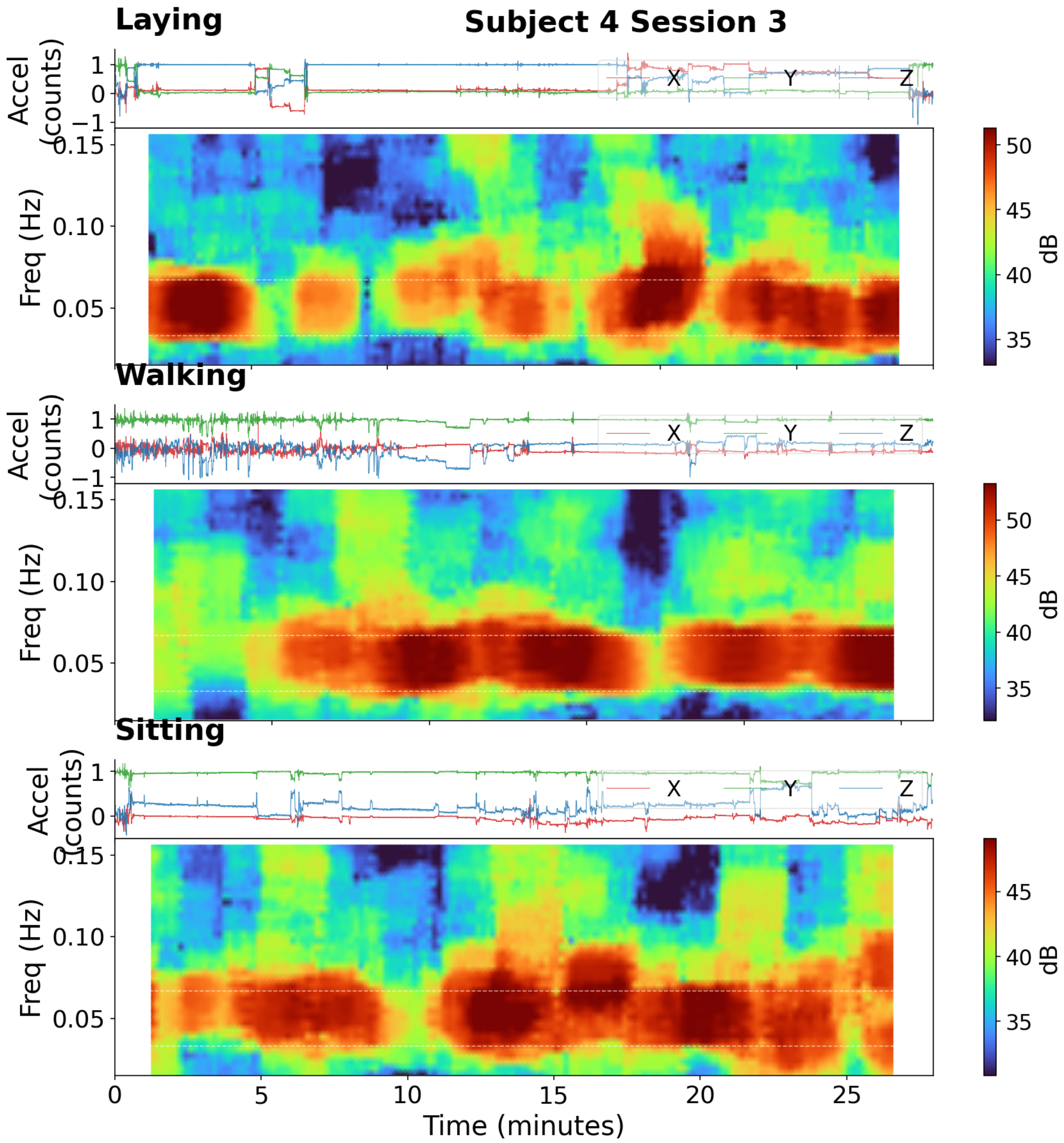


Figure S38. (Subject S4). Post-meal activity phases: laying, walking, sitting. Top panel: triaxial accelerometry. Bottom panel: multitaper HR-EGG spectrogram (120-s window, 15-s step). Dashed white lines mark the 2–4 cpm normogastric band.

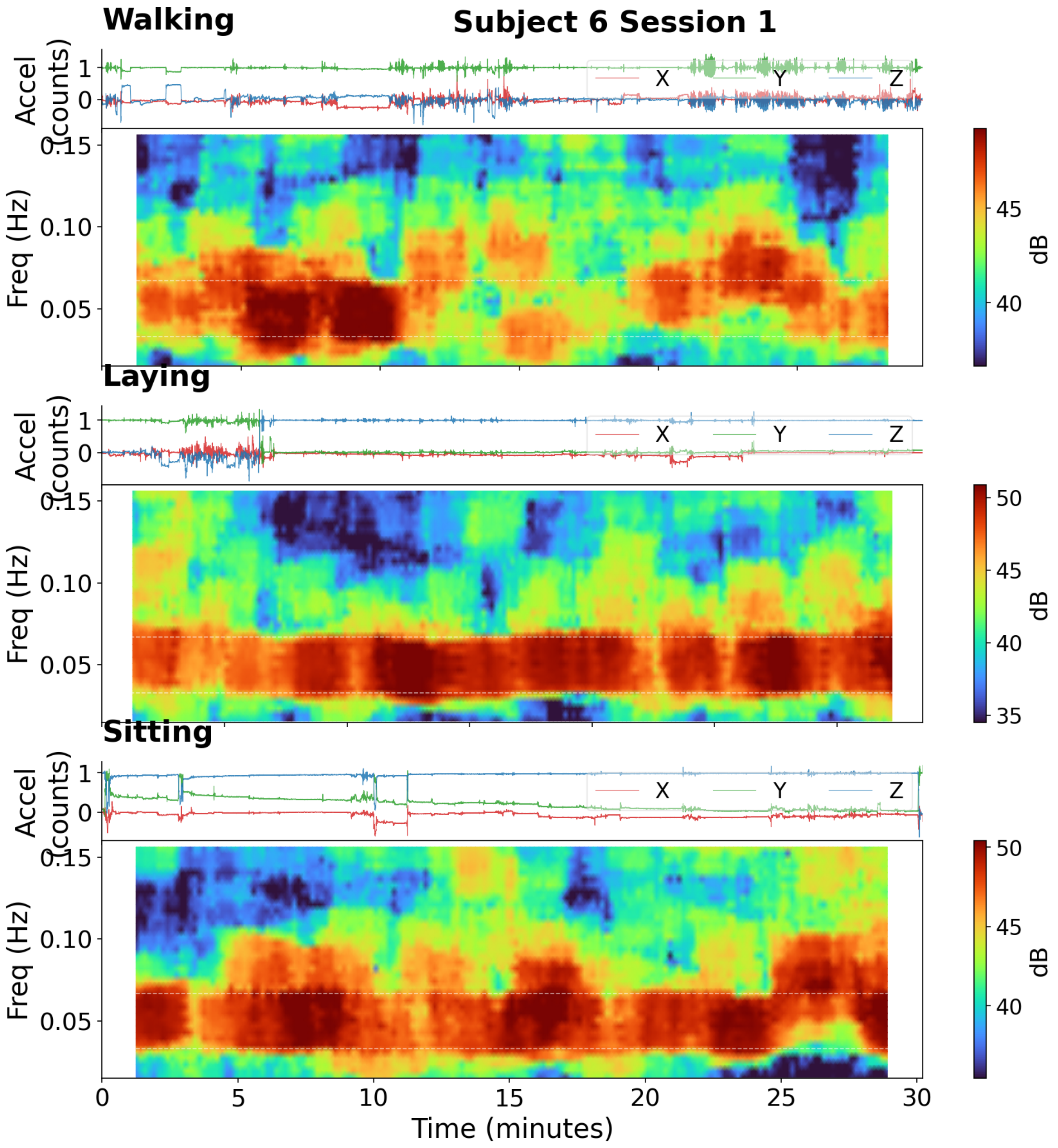


Figure S39. (Subject S6). Post-meal activity phases: walking, laying, sitting. Top panel: triaxial accelerometry. Bottom panel: multitaper HR-EGG spectrogram (120-s window, 15-s step). Dashed white lines mark the 2–4 cpm normogastric band.
